# Shallow-water chemosymbiotic clams are a globally significant and previously overlooked carbon sink

**DOI:** 10.1101/2024.02.25.581922

**Authors:** Menggong Li, Yunlong Li, Shi-Hai Mao, Zhixin Zhang, Chong Chen, Xueying Nie, Xu Liu, Hui Wang, Xiaoshou Liu, Weipeng Zhang, Qiang Lin, Guang-Chao Zhuang, Jin Sun

## Abstract

Chemosynthetic animal symbioses are common in marine ecosystems but remain overlooked as contributors to global carbon fixation. We show that the shallow-water thyasirid clam *Thyasira tokunagai*, dominant in Yellow Sea sediments, harbors sulfur-oxidizing *Sedimenticola* symbionts with remarkably consistent genomic contents and functionality across the region, showing active Calvin cycle gene expression and close-knit host-symbiont metabolic integration. Field surveys demonstrated densities up to 2015 individuals·m□^2^, while radiocarbon tracing revealed assimilation rate constants (0.002–0.005 day□¹) peaking at 14.8°C. Spatial modelling combining abundance and temperature estimated a carbon fixation of 0.89 Tg C·yr□¹ in Yellow Sea, equivalent to 43% of the annual sedimental C burial from the Chinese coast. The species complex that includes *T. tokunagai* is widely distributed and constitutes a globally significant, previously unaccounted blue carbon sink. Our findings underscore the crucial role of shallow-water chemosymbioses in carbon cycling, emphasising the importance of incorporating them into climate models and conservation strategies focused on carbon sequestration.

## INTRODUCTION

Carbon fixation is a fundamental process converting inorganic carbon into organic compounds and underpins the entire global food web (*1*). Photosynthesis, a light-driven process, accounts for the majority of primary productivity on Earth. Chemosynthesis, best known from deep-sea environments, represents the more ancient form of primary production. Chemoautotrophic microorganisms obtain the chemical energy to fix inorganic carbon through the oxidation of reducing substances, such as sulfides, methane, and hydrogen gas, providing an alternative carbon fixation pathway to photosynthesis (*2*). Increasing evidence indicates that chemosynthesis supports diverse microbial and animal life across the entire ocean, influencing the global biogeochemical cycling of nutrients and the marine carbon budget (*2–4*).

Chemosynthetic microorganisms are widely distributed throughout marine ecosystems, inhabiting all depth zones and latitudinal gradients (*2*). Most studies on dark carbon fixation in marine ecosystems focus on free-living chemoautotrophic bacteria. Significant dark carbon fixation also occurs in coastal sediments, where maximum chemoautotrophy rates reach 3-36 mmol C·m^−2^·d^−1^ in the upper 1-2 cm (*5*). In the deep ocean, benthic dark inorganic carbon fixation (DCF) rates ranges between 1.51*10^1^-3.24*10^2^ μg C·m^−3^·h^−1^ in the water column and 1.15*10^4^-1.83*10^5^ μg C·m^−3^·h^−1^ in sediments (*3*). In contrast to free-living chemoautotrophic bacteria, chemosymbiotic animals house chemosynthetic microbes in specific organs or cells, functioning as ‘ecological containers’ (*6*). This intimate partnership is exemplified by symbiotic tubeworms, bivalves, and gastropods inhabiting deep-sea vent and seep ecosystems (*6–8*). Chemosymbiosis also extends to shallow-water environments, however, encompassing ciliated protists, nematodes, gutless oligochaetes, and lucinid clams associated with seagrass or mangrove sediments, as well as thyasirid bivalves inhabiting organic-rich, cold-water sediments (*8*). Biological carbon fixation exerts a notable influence on global carbon budgets (*9*). Despite the recognized importance of chemosymbiosis, the contribution of shallow-water chemosynthetic symbioses to regional or global carbon budgets remains unquantified. So far, only one study estimated the carbon and nitrogen fate in shallow-water lucinid clams, using stable isotope incubation – with an estimated carbon fixation rate of 300-2600 nmol C·g gill tissue^−1^·h^−1^ – but less is known about the carbon fixation budget (*10*).

Here, we use a thyasirid clam prevalent in the reducing sediments in the Yellow Sea – *Thyasira tokunagai* (**Fig. 1A**) *–* as a model for quantifying the carbon fixation contribution of chemosymbiotic animals in shallow water. *Thyasira tokunagai* is a member of the widely distributed *T. gouldii* species complex commonly found in reducing sediments all around cold shallow-water habitats around the northern hemisphere from eastern Canada to Europe to Asia (*11–13*). Quantifying the carbon fixation contribution of the *T. tokunagai* is crucial for a more complete understanding of carbon cycling in the Yellow Sea and other similar coastal ecosystems (*14*), which provides an opportunity to address this gap. Our findings highlight chemosymbiotic animals as a previously unappreciated carbon sink in shallow water, comparable to well-known carbon sequestration ecosystems such as mangroves and seagrass beds.

**Fig. 1.**
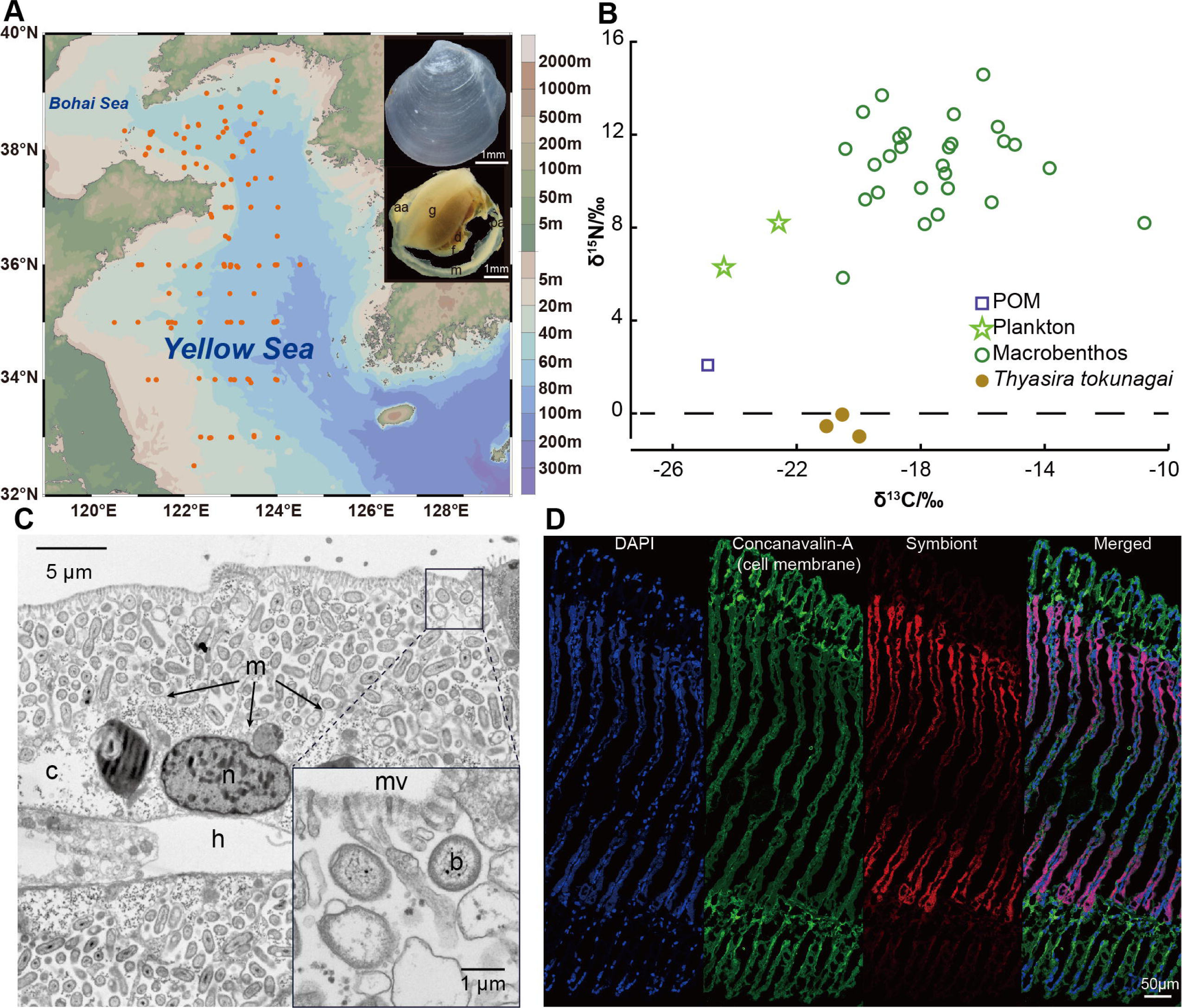
Sampling sites, stable isotope signatures, and chemosymbionts. **(A)** Shell and external anatomy of *T. tokunagai*, and a total of 139 sampling sites during seven research cruises from 2018-2024 in the Yellow Sea. Abbreviation: g, gill; f, foot; d, digestive diverticula; m, mantle; aa, anterior adductor; pa: posterior adductor. **(B)** The stable isotopic niche of *T. tokunagai* in the macrobenthic community of the Yellow Sea. Additional data on other macrobenthos in the Yellow Sea can be found in Table S2 (*15*). The red dots represent *T. tokunagai.* **(C)** Transmission electron microscopy (TEM) image of extracellular bacteria maintained in pouch-like structure bearing microvilli. Abbreviation: mv, microvilli; b, bacteria (the high abundance bacteria are mainly symbiont); n, nuclei; m, cell membrane; c, cell cytoplasm; h, hemocoel. **(D)** Fluorescence *in situ* hybridization (FISH) showing the symbiont bacteria in the gill filaments. All cell nuclei were stained with DAPI, the cell membrane was stained by concanavalin-A, and symbionts were hybridized by the specific probe of the *T. tokunagai* symbiont *Sedimenticola* sp.

## RESULTS

### Host-Symbiont System in a Homogeneous, Shallow Environment

Our sampling records across seven cruises between 2018 to 2024 show that *Thyasira tokunagai* is a dominant benthic species in the Yellow Sea, with up to 2015 individuals per square meter, typically found at depths ranging from 9-82 meters **(table S1)**. *Thyasira tokunagai* was identified from both morphology and the mitochondrial *cox1* gene barcode to confirm its placement in the *Thyasira gouldii* species complex (**Fig. 1A and fig. S1A**; personal communications, Suzanne Dufour). Pairwise comparisons of the *cox1* gene of specimens collected from nine sampling locations in the Yellow Sea revealed an average of 99.84% similarity (**fig. S1B**), and the haplotype network (**fig. S1C**) also showed that nearly all haplotypes lacked a clear geographical affinity. Furthermore, our STRUCTURE analysis also showed a lack of population differentiations using the alignment of 13 protein-coding genes (PCGs) of 30 mitochondrial genomes based on the delta *K* (*K* = 2; **fig. S1D**). These results indicate a panmixia condition for all nine populations of *T. tokunagai* sampled in the Yellow Sea.

Compared to other benthic fauna and environmental samples from the Yellow Sea (*15*), *T. tokunagai* exhibited the lowest δ^15^N value (−0.23 ± 0.22 ‰, n = 3), suggesting a potential autotrophic supplement on its nitrogen source (**Fig. 1B and table S2**; particulate organic matter (POM): 2.08 ‰, phytoplankton: 6.28 ‰, benthic fauna: 10.77 ‰). The δ^13^C value of *T. tokunagai* (−20.52 ± 0.43 ‰, n = 3) was higher than that of the POM and phytoplankton, but lower than other benthic fauna (POM: −24.87 ‰, phytoplankton: −24.35 ‰, benthic fauna: −17.46 ‰). These results indicate that *T. tokunagai* does not rely on the filtration or predation of other organisms for nutrition, though it may obtain organic matter from others.

A bacterial species belonging to the genus *Sedimenticola* dominated the bacterial community in the gills of *T. tokunagai* (**fig. S2**), determined through full-length 16S rRNA gene amplicon sequencing, showing this *Sedimenticola* species accounted for an average of 92.34% of the bacterial community (n = 21). Phylogenetic reconstruction using the 16S rRNA gene confirmed that its closest relative was the chemosymbiont of *T.* cf. *gouldii* in the same species complex (**fig. S3**). To investigate the distribution of symbionts within the gill tissue, we conducted fluorescent *in situ* hybridization (FISH) imaging, which show that 1) the symbionts were concentrated in the bacteriocytes located at the middle part of the gill filament except the ciliated filament tip (**Fig. 1C and fig. S4**) and 2) symbionts seemed to be enveloped by a layer of membrane (indicated by the green signal). Nonetheless, transmission electron microscopy (TEM) observation (**Fig. 1D**) of the gill tissue showed that symbionts were actually localized in extracellular pouch-like structures among the microvilli but not completely enclosed in vesicles, implying an exocellular symbiotic mode where bacteria are maintained outside of the host cytoplasm but in a specialized pouch-like organ (*16*).

### Two Symbiont Phylotypes

Two phylotypes of the same *Sedimenticola* species made up the bacterial population in *T. tokunagai*. There was only a single base pair difference (G vs A) between these two phylotypes at the 590^rd^ position of the 16S rDNA, verified by Sanger sequencing (**Supplementary Note 1**) – each phylotype accounting for 46.29% and 45.88% of the overall symbiont population (mean percentage), respectively (**Fig. 2A**). Host individuals differ greatly in the proportion of the two phylotypes, showing a whole range including some with only one or the other phylotype (**Fig. 2B**). Spot-analyses (10 µm in diameter) of gills showed that over 80.59% of 10,468 spots exhibited just one phylotype, indicating there is a bias to hosting just one phylotypes within each symbiotic pouch. H&E imaging and FISH analysis confirmed the symbiont distribution pattern on the gill filament (**Fig. 2, C and D**). Spatial analyses of gills show that individuals also vary in the level of spatial heterogeneity concerning the phylotypes (two gills per individual, **Fig. 2E**) – especially in the individual g5 (a total of six individuals), which showed well-mixed patterns of the two phylotypes.

**Fig. 2.**
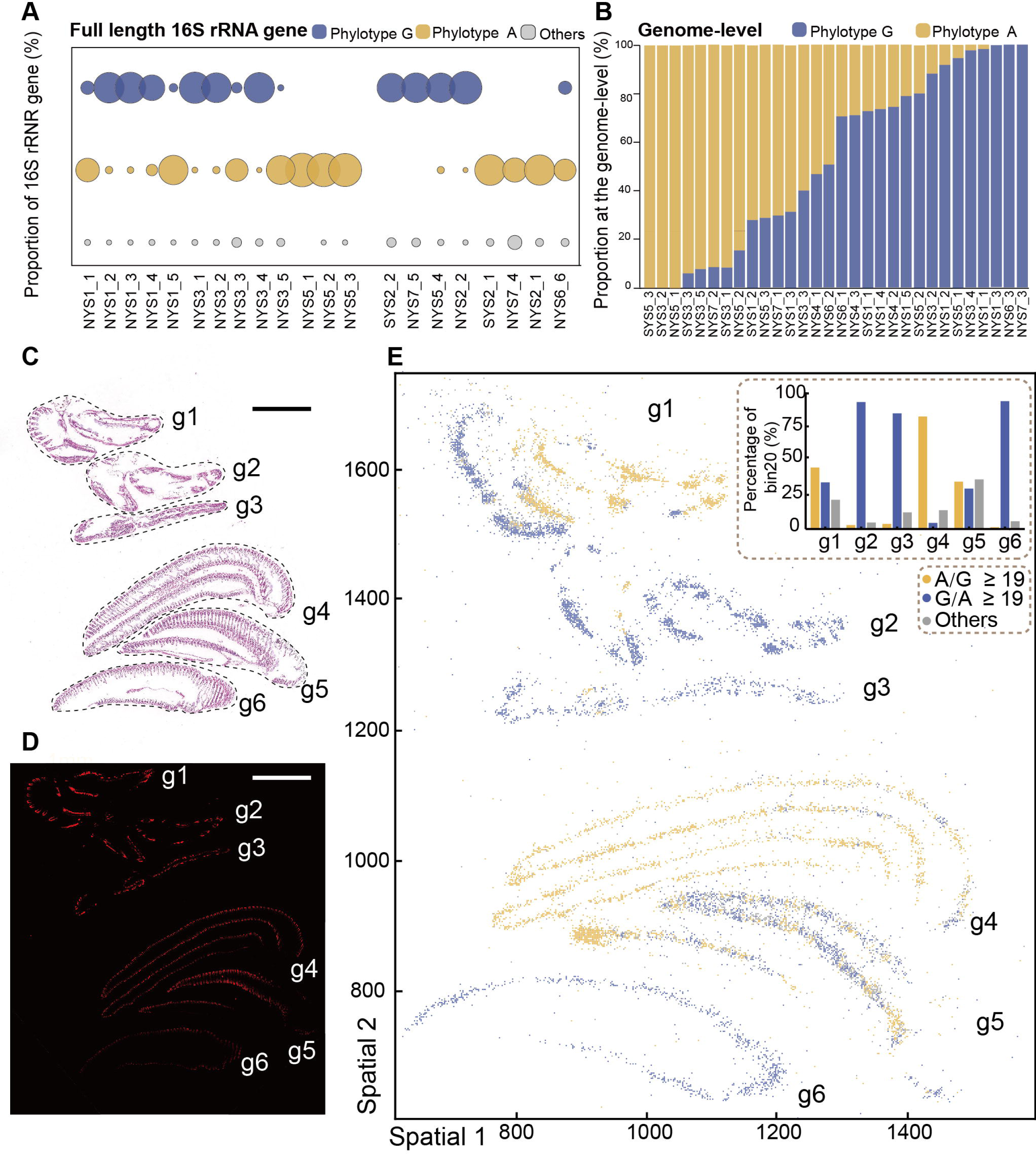
Identification and spatial distribution of the two *Sedimenticola* symbiont phylotypes in *Thyasira tokunagai* gill tissue. **(A)** Two dominant phylotypes belonging to the genus *Sedimenticola* were identified in gill tissue bacterial communities which we call *Sedimenticola* sp. (ex *Thyasira tokunagai*) ‘phylotype A’ and ‘phylotype G’ based on 16S rRNA gene sequencing, differing by a single base at the 563rd position; the first 13 samples were used for metagenomic sequencing, while the last 8 were used for metatranscriptomic sequencing. NYS represents the sampling sites from the north Yellow Sea, whereas SYS represents the sampling sites from the south Yellow Sea. **(B)** Strain decomposition analysis revealed the presence of two symbiont phylotype, corresponding to the two dominant phylotypes identified in (A). **(C)** The H&E-stained gill tissue section (scale bar: 1 mm) had a neighboring section used for spatial phylotype distribution analysis (see panel E). **(D)** FISH imaging showing the symbiont-specific probe visualized spatial distribution in a gill tissue section (scale bar: 1 mm), while a neighboring section was observed after hematoxylin and eosin (H&E) staining (see panel C and E). **(E)** At the cellular level, the spatial distribution of symbiont phylotypes is such that each spot represents a 10 μm bin, named bin 20, approximating the size of a single cell, and the A/G ratio indicates the relative abundance of phylotype A compared to phylotype G at each location (e.g., A/G > 19 indicates that phylotype A is approximately 19 times more abundant than phylotype G within that bin).

Combining long-read and short-read sequencing, a total of 30 high-quality circular genomes (i.e. MAGs) were assembled, with completeness > 99.23%, contamination rate < 0.34%, and size of 4.5 Mb (**Fig. 3A and table S3**). The average nucleotide identity (ANI) of these 30 MAGs ranges from 98.90 to 99.94% (**fig. S5 and table S4)**, above the proposed threshold of inter-species variation of prokaryotes (95%) and supports them as belonging to the same species (*17*). Representative genome of each phylotype was further deduced using StrainPanDA (**Fig. 3A)**. Genome capacity analysis revealed no functional gene differences between the two phylotypes **(table S5).** To boost the credibility of our findings, we conducted a correlation analysis. This analysis revealed a positive correlation between the percentage of phylotype G from StrainPanDA and the percentage of metagenomic reads of G base pair at the 563^rd^ position in 16S rRNA gene (**fig. S6; *R*^2^ = 0.97, *P* < 0.001**). Their placement in the genus *Sedimenticola* was also shown by phylogenetic reconstruction at the genomic level (**Fig. 3B**). Overall, it seems the two phylotypes are equivalent in function and the host individuals do not actively select for one or the other phylotype, instead using them interchangeably.

**Fig. 3.**
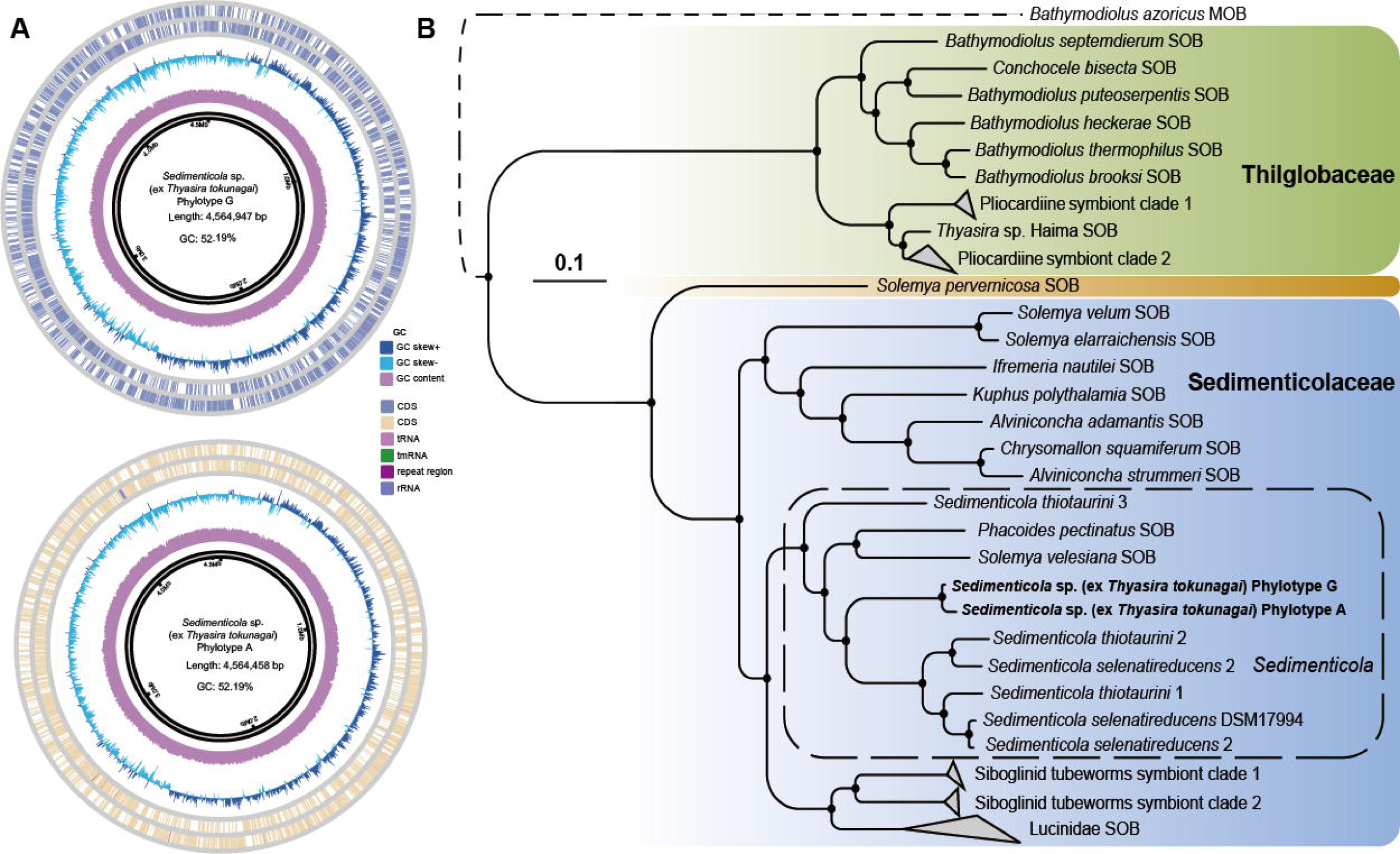
Phylogenetic relationships of the two *Sedimenticola* symbiont phylotypes. **(A)** General features of the symbiont genomes showing that the two genomes have a similar size (a total of 4.5 Mb) and a GC content of 52.19%. **(B)** Phylogenomic analysis, rooted with the methanotrophic (MOB) symbiont of the hot vent bathymodioline mussel *Bathymodiolus azoricus,* demonstrates that the two phylotypes fall within the family Sedimenticolaceae (SOB) and are closely related to cultured *Sedimenticola* species (solid black dot at the node indicates 100 bootstrap supports; scale bar: 0.1 substitutions per nucleotide site).

### Symbiont Chemosynthetic Capacity Ensures Carbon Fixation

The metabolic potential of the *Sedimenticola* symbiont of *T. tokunagai* was highly conserved (**Fig. 4A**), highly similar to the published result in the closely related *T.* cf. *gouldii* (*18*). They encode the full set of enzymes in carbon fixation and utilization, including the Calvin–Benson–Bassham cycle (CBB cycle, or the reductive pentose phosphate cycle), glycolysis/gluconeogenesis, tricarboxylic acid cycle (TCA), and oxidative phosphorylation, enabling both phylotypes to assimilate dissolved inorganic carbon. The ribulose-bisphosphate carboxylase large chain (*rbcL*, K01601) in the Calvin cycle is responsible for the assimilation of inorganic carbon. The reductive TCA cycle cannot function completely in *T. tokunagai* symbiont due to the absence of type II ATP citrate lyase, unlike the symbiont of the giant tubeworm *Riftia pachyptila* (*19*). The complete dissimilatory nitrate reduction pathway proceeds with respiration under an anaerobic or hypoxic environment but hinders it from producing ammonia due to the lack of the nitrite reductase (NADH) large subunit (K00362). A complete dissimilatory sulfate reduction pathway and a partial SOX system were also found, mainly containing *soxA*, *B*, *X*, *Y*, and, *Z*, but *soxCD* was lacking. The incomplete assimilatory sulfate reduction pathway was detected, which contained *sat* and *cysC*. We also found the genes and enzymes related to hydrogen oxidation, containing *hoxF*, *U*, *Y*, *H*, *hybC*, and *hyaB*. The symbiont encodes ABC transporters and PTS pathways, indicating the capacity of heterotrophy. The bacterial chemotaxis and flagellar assembly pathways were found. Additionally, both phylotypes have a relatively complete capacity for the biosynthesis capacity of amino acids (17), vitamins, and cofactors (10) **(table S6)**, suggesting a capacity for free-living habit. The genes in the pathways that are highly relevant to chemosynthesis are expressed actively, such as the CBB cycle, sulfur oxidation, and nitrogen metabolism (**Fig. 4B**).

**Fig. 4.**
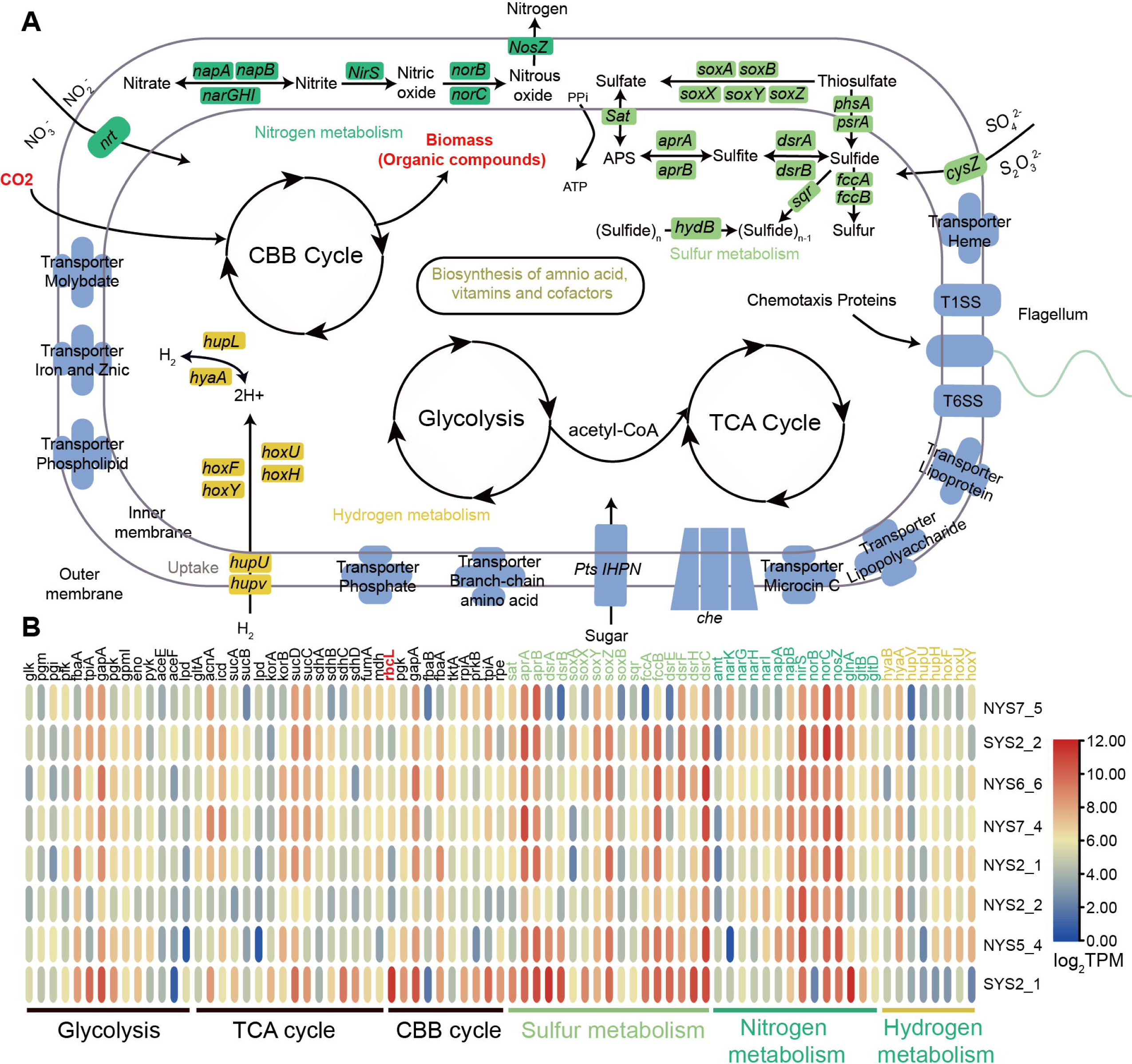
Chemosynthetic capacities and transcriptome profiles of symbiont phylotypes. **(A)** Reconstructed chemosynthesis pathways from both symbiont genomes indicate a conserved metabolic potential, consistent with a free-living lifestyle. Both phylotypes encode a complete set of enzymes for carbon fixation and utilization and have the potential to utilize sulfides and hydrogen as energy sources. **(B)** Transcriptome profiles of highly conserved chemosynthetic pathways related to carbon, sulfur, nitrogen, and hydrogen metabolism. Transcribed functional genes were annotated against the KEGG database, and expression levels are shown as log2-normalized Transcripts Per Kilobase Million (TPM).

### Carbon fixation flux estimation

Since the *Thyasira gouldii* complex is widely distributed and abundant across shallow cold-water habitats of the northern hemisphere, with potential to be a carbon sink, we proceeded to estimate the carbon fixation flux and carbon sequestration capacity of *T. tokunagai* as a model (**Fig. 5A**). The enzyme activity in animals alters with temperature. To assess the carbon assimilation activity of clams, we added ^14^C-labeled DIC tracers to homogenized bacterial solution from the same sampling site (four sites in total in this study, **Fig. 5B**). Considering the impact of varying symbiont numbers, we chose four replicate samples from each site. All samples for each site were incubated at four temperatures (i.e., 5, 12, 20, and 28□°C), covering the lowest and highest levels across a whole year in the natural habitat (**Fig. 5C and table S7**). Results showed that the carbon assimilation rate constant (*k*) was lowest at 5□°C across all sites (average: 0.002148 ± 0.001372). As temperature increased, the *k* exhibited higher levels, with the highest and comparable values at 12 and 20 °C (0.005092 ± 0.000448 and 0.004788 ± 0.000976, respectively). A declining trend was observed with the continuously rising temperature until 28 °C (*k =* 0.003508 ± 0.001204), only accounting for 69% of the highest *k* value. The Wilcoxon Signed-Rank test (**fig. S7**) indicated significant differences at the same temperature, but only between some sites, suggesting that symbiont copy number and metabolic activity may influence carbon assimilation activity. Nonetheless, symbionts in *T. tokunagai* appear to transfer more DIC to organic matters at elevated temperature. A cubic regression model presented a better performance in regression from measured value, than a quadratic one (**Fig. 5C and fig. S8**), predicting a peak assimilation rate constant at 14.78 °C, with decreasing under both warmer and cooler conditions.

**Fig. 5.**
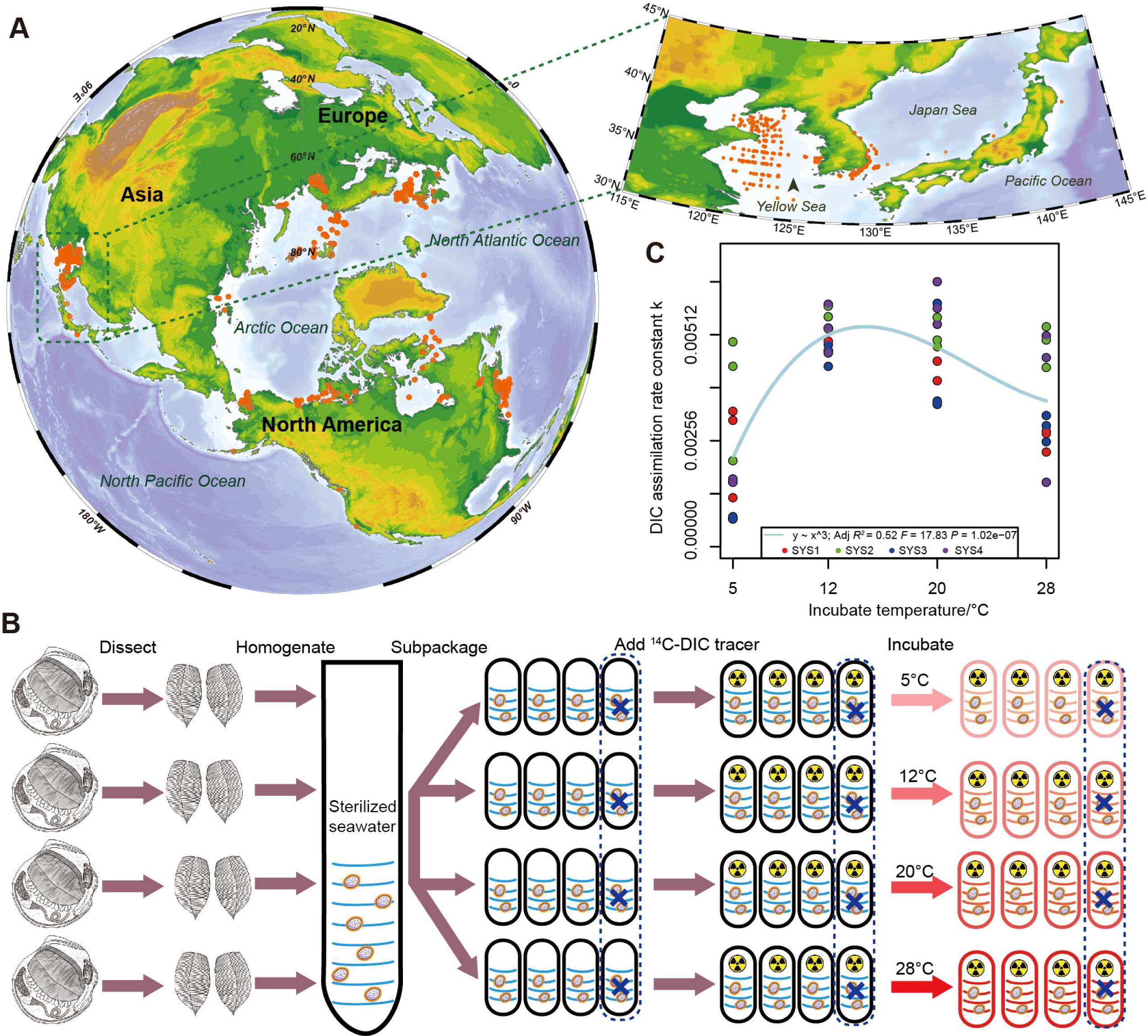
Global distribution of the *Thyasira gouldii* complex and the carbon fixation rate of *T. tokunagai*. **(A)** Global distribution of the *Thyasira gouldii* complex, and *Thyasira tokunagai* distributed in the Yellow Sea and Japan Sea. **(B)** Flow of the ^14^C-labeled DIC assimilation assay. To assess DIC assimilation, gill tissue samples were dissected, homogenized in sterilized seawater, and aliquoted into sealed penicillin bottles. ^14^C-labeled DIC tracer was then added, and the samples were incubated for nearly two days at various temperatures (5, 12, 20, and 28 □) before measuring carbon assimilation rates. The symbol of “X” represents the negative control treated by trichloroacetic acid. **(C)** Carbon fixation rate constants at different temperatures (5°C, 12°C, 20°C, 28°C) fitted using a trinomial equation.

Previous cruises and research reported that the dissolved inorganic carbon (DIC) in the bottom water fluctuated within a small range between 2.03 and 2.23 mmol/L (*20*). Furthermore, we applied kriging interpolation to estimate the annual average bottom-water temperature (average level in year) and the spatial distribution of *T. tokunagai* across the Yellow Sea (**Fig. 6, A and B**). The results showed that annual bottom-water temperature ranged from 6 to 16 □, and three major inhabiting groups aggregated in the active cold-water mass regions (**Fig. 6, A and B**). Based on the established relationship between temperature and the carbon assimilation rate constant (**Fig. 5C**; *R^2^* = 0.52, *P* = 1.02e^−07^), we estimated the spatially resolved carbon fixation rate of *T. tokunagai* symbionts in the Yellow Sea (**Fig. 6C**). The cold-water mass of northern region in Yellow Sea was the central contributor in carbon assimilation, with a peak fixation rate of 27.1791 g·m^−2^·yr^−1^, coinciding with areas of high *T. tokunagai* abundance. Considering the areal extent of the Yellow Sea, we estimated that the total annual carbon fixation rate by *T. tokunagai* symbionts in the region was approximately 0.892 Tg·C·yr^−1^.

**Fig. 6.**
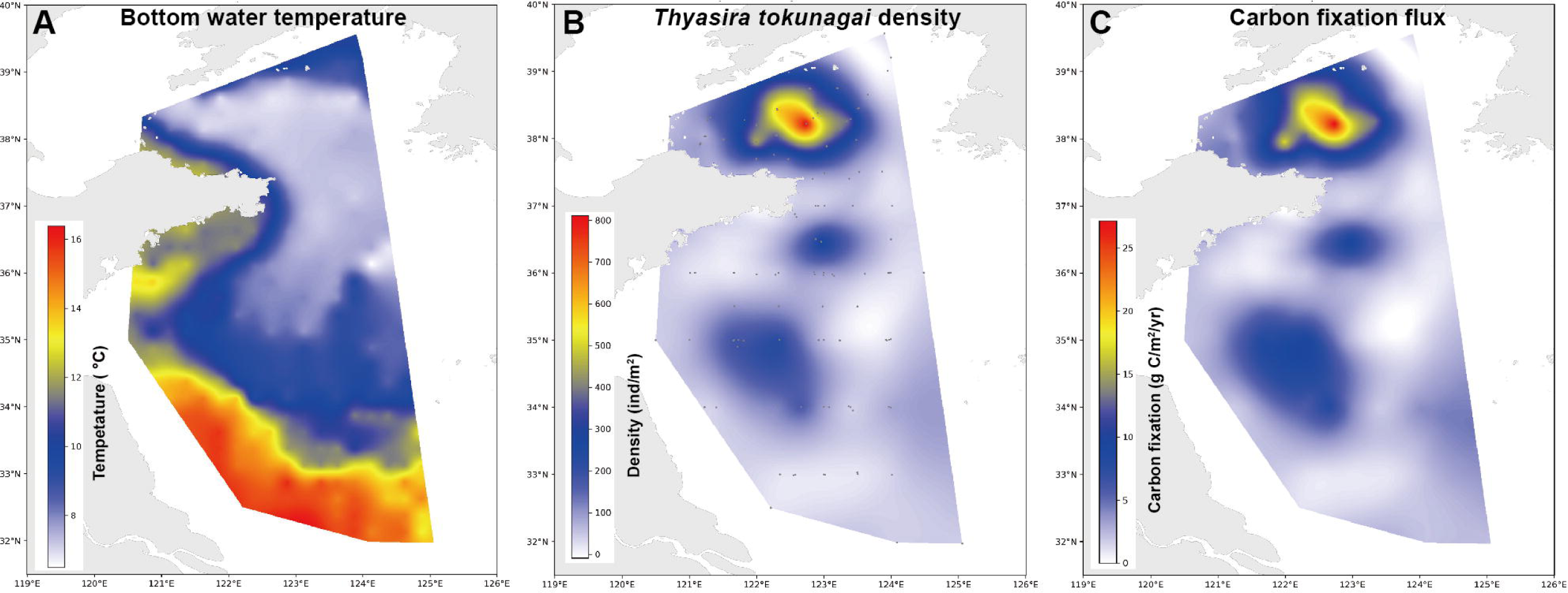
Estimation of the total carbon fixation flux of *Thyasira tokunagai* in Yellow Sea. **(A)** Visualization of the mean annual bottom water temperature from 2002-2017, and this data was used to predict carbon fixation rate constants. **(B)** *T. tokunagai* density in the Yellow Sea. Based on data from 139 sites across seven cruises, we predicted density using the kriging interpolation method within the sampling area for carbon fixation flux estimation. **(C)** Annual carbon fixation flux within the sampling area is estimated using the kriging interpolation method, units represent the total annual fixed carbon per site.

## DISCUSSION

### Chemosynthetic capacity of thyasirids in the Yellow Sea

Chemosymbiosis is widely distributed in Mollusca, but most studies have been undertaken in deep-sea vents and seeps (*6, 7*). Most members of the family Thyasiridae are widely distributed in non-vent/seep anoxic mud, including shallow-water ecosystems, but they remain little-studied and their role in ecosystems much overlooked. Our results are the first detailed study on the symbiosis of *Thyasira tokunagai* despite its abundance, revealing the dominance of a sulfur-oxidizing bacterial (SOB) symbiont belonging to *Sedimenticola* in their gills clustering closely with other previously reported chemosymbionts of other animals (**fig. S2**).

Consistent with previous metagenomic work on the closely related congener *T.* cf. *gouldii* (*18*), the *T. tokunagai* symbiont genomes also exhibited *soxA, B, X, Y,* and *Z*, but not *soxCD*. The absence of *soxCD* would lead to the partial oxidation of thiosulfate and the accumulation of zero-valent sulfide, forming sulfur globules in the symbionts, which are common in SOB such as *Erythrobacter flavus* (*21*) and *Chlorobaculum limnaeum* (*22*). Sulfur globule-like structures were observed in the symbiont of both *T. flexuosa* and *T.* cf. *gouldii* under TEM (*23, 24*). On the other hand, genomes of our *T. tokunagai* symbiont contained a complete dissimilatory sulfate reduction (*dsr*) pathway, suggesting the genetic basis to fully oxidize sulfide to sulfate (*25*). During that process, ATP is generated by sulfate adenylyltransferase (*sat*) and then utilized in the Calvin cycle to fix CO_2_, which is a central principle of chemosymbiosis. Additionally, *T. tokunagai* symbionts appear to be capable of fixing inorganic carbon via oxidizing reducing substances, with the SOX system, reverse dissimilatory sulfate reduction pathway, and the HOX system (*26, 27*). The capability of dissolved inorganic carbon assimilation has been quantified, providing robust support of hosting chemosymbiont in *T. tokunagai*.

### Strong selection of host on symbiont and horizontal transmission

Though our sampling sites of *T. tokunagai* covered a wide area in the Yellow Sea, with the longest distance from the southernmost point to the northernmost point over 500 km, there was an undifferentiated population panmictic across all sites as evidenced by mitochondrial population genomics. The presence of the symbiont population in the gill of *T. tokunagai* is greatly dominated by a single symbiont species (genus *Sedimenticola*) points to the strong selection of symbionts by the host (*28*). The ratio of the two phylotypes within each host likely result from local bias within each symbiont-hosting pouch (spatial metabarcoding result in **Fig. 2E**). Overall, we observed a lack of bias in phylotype uptake by the host, with a comparable percentage of the two phylotypes across the whole *T. tokunagai* populations and a spatial section consisting of the mixed pattern. The two phylotypes differ only by one single base pair in the 16S rRNA gene and appear to be without functional differentiation, supported by the highly similar gene sets between the two deduced phylotypes’ genomes. This is different from the symbiont phylotypes diversity in deep-sea bathymodioline mussels, which has a significant impact on the fitness (*29*). Our results suggest all individuals of *T. tokunagai* share symbionts of a consistent function and ecology – enabling a reliable extrapolation of our measured carbon fixation rate to the whole Yellow Sea population.

Previous studies reported the super extensile foot in thyasirid bivalves to mine sulfide and the magnetosome in its symbionts (*30, 31*), implying the symbionts might be derived from the specific niche of sediments. The metabolic potential of the symbiont genomes, with the capacity for heterotrophy and free-living, is suggestive of horizontal transmission. Therefore, we consider *T. tokunagai* most likely acquire the *Sedimenticola* symbionts from the environment via horizontal transmission, acquired under the control of a highly selective mechanism.

### An overlooked sink of carbon dioxide from chemosynthetic thyasirids in shallow water

Global climate change, primarily driven by anthropogenic activities, is occurring at an unprecedented pace compared to past climatic fluctuations (*32, 33*). Achieving net-zero CO_2_ emissions is critical to mitigating this trend, necessitating accurate quantification of carbon sinks, particularly blue carbon, i.e., carbon captured by marine ecosystems. Previous research has largely focused on conventional blue carbon contributors such as phytoplankton, marine microbes, mangroves, and seagrasses (*9, 34*). However, holobiont animals that fix carbon through chemosynthesis primarily using the CBB cycle with minor contribution from the rTCA cycle may represent an underestimated and significant carbon sink. Compared to charismatic species such as the giant tubeworm *Riftia pachyptila* that only live-in biodiversity hotspots like vents, chemosymbiotic bivalves in families like Thyasiridae and Lucinidae in anoxic shallow-water habitats are distributed across vast geographic ranges and should represent a much higher total carbon flux. Despite this, few studies have explored the rate of carbon assimilation in chemosynthetic holobionts and their potential importance for oceanic carbon fixation.

Here, we combine genomic and transcriptomic analyses and reveal the molecular mechanisms underlying carbon fixation in the chemosymbionts of *Thyasira tokunagai*. Using radiocarbon tracing technique, we also quantified its carbon fixation rate. Our results revealed that each *T. tokunagai* individual can fix up to 34.3–37.9 mg C·yr^−1^, with rates mainly affected by symbiont density and the ambient temperature. Nevertheless, based on the population density of *T. tokunagai*, we approximated an aerial carbon fixation rate of up to 27.2 g C m^−2^ yr^−1^ based on our rate measurement. Importantly, our multi-omics analyses show that the symbiont populations of *T. tokunagai* across the entire Yellow Sea is remarkably consistent in genomic content with only two phylotypes that share the same functionality; meaning our estimated rate can be reliably applied to all individuals of *T. tokunagai*. This value is significantly higher than the carbon fixation rates reported for many shallow-water and terrestrial ecosystems, as well as deep-sea dark DIC fixation rates, such as lake sediment, seawater, and soil (*35, 36*). Although the rate is lower than those of mangroves (170-190 g C·m^−2^·yr^−1^) and seagrass (5-379 g C·m^−2^·yr^−1^), the broad distribution and high abundance of *T. tokunagai* in the Yellow Sea compensates it in scale, yielding an estimated total carbon fixation of 0.892 Tg C·yr^−1^. This value is comparable to 43% of the total carbon burial rates along the Chinese coast (up to 2.06 Tg C·yr^−1^), which consisted of 0.05 Tg C·yr^−1^ from mangrove (25,900 ha), 0.50 Tg C·yr^−1^ from salt marsh (297,900 ha), and 0.28-1.5 Tg C·yr^−1^ from tidal flat (237,450-1,102,400 ha) (*34*). If counted based on the capability of mangrove (~ 180 g C·m^−2^·yr^−1^ averagely), the contribution from *T. tokunagai* in the Yellow Sea is equivalent to about 495,000 ha mangrove and 2% of global mangrove (*34*). Furthermore, thyasirids burrow deep in the anoxic mud and may tend to reserve the fixed carbon in the sediment, based on moderated turbulence at the sediment-water interface compared to the more dramatic waves in the intertidal region where mangroves and seagrasses are located.

The *Thyasira gouldii* complex, which *T. tokunagai* is a part of, is widely distributed across the cold-water region across the entire northern hemisphere (**Fig. 5A**). Like the symbionts of *T. tokunagai*, those of *Thyasira* cf. *gouldii* have a complete Calvin–Benson–Bassham (CBB) cycle, indicating a high carbon fixation potential for the *Thyasira gouldii* complex (*18*). We found a temperature dependence of carbon assimilation in chemosymbiotic thyasirids, while the typical bottom-water temperature in Yellow Sea where the majority of *T. tokunagai* inhabits is between 6-10 °C, within the climbing curve of the rate constant-temperature relationship (**Fig. 5C**). Recently, marine heatwaves have been increasingly observed in the deeper layers, persisting for longer and more intense than the surface seawater (*37*). For instance, bottom water temperatures at depths of 50–100 m along the North American continental shelf increased by up to 3□°C between 1993 and 2019, with similar phenomenon observed in the deep water of high latitude regions (*38, 39*). Previous projections had indicated an increase of 0.5 °C in the cold water mass by the end of 2050 (*40*), which could enhance carbon fixation by 4.37% by the chemosymbionts of *T. tokunagai* by the end of 2050, if the survival and distribution of *T. tokunagai* are not significantly affected by this (**fig. S9A**). Likewise, a further increase of 2 °C by 2100 could increase carbon fixation by up to 17.71% (**fig. S9B**). These predictions highlight the potential increase of carbon fixation by shallow-water chemosymbiotic communities in a climate change scenario.

Collectively, our study used multi-omics and radiotracer techniques to elucidate the carbon fixation potential of shallow-water thyasirid holobionts. Our findings reveal a previously unrecognized carbon sink and highlight the ecological importance of shallow-water chemosynthetic symbioses in marine carbon cycling. Considering that *T. gouldii* complex (including *T. tokunagai*) is a dominant species in the Yellow Sea, Japan Sea, as well as the pan-Arctic area in Atlantic and Pacific Ocean (*11, 12*), the *T. gouldii* complex or other chemosymbiotic species may play a more substantial role in marine DIC fixation than previously recognized. Further ecological investigation into the distribution and density of these chemosymbiotic organisms, along with in-depth studies of the factors influencing their carbon fixation rate, is crucial to accurately assess their contribution to global carbon cycling.

## MATERIALS AND METHODS

### Sampling description

*Thyasira tokunagai* (**Fig. 1A**) were collected from a total of 139 sites in the Yellow Sea between 42-72 m depth from seven cruises on-board the R/V *Lanhai 101* from 2018 to 2024, with the site details and environmental parameters shown in **table S1**. A 0.1 m^2^ box corer was employed to collect the surface sediments. The *T. tokunagai* were manually picked out from the sediments via a 0.5 mm sieve once they were on board and then immediately fixed and preserved. For details on the sample preservation, please see the supplementary information.

### Stable isotope analysis

Stable isotope analysis of carbon (C) and nitrogen (N) was conducted as previously described (*41*). The whole tissues of three *T. tokunagai* specimens were freeze-dried for two hours at −60□, and approximately 0.1 mg of the powdered sample was analyzed for stable isotopes using an isotope ratio mass spectrometer (IRMS, Sercon Instruments, Crewe, UK) at the Third Institute of Oceanography, China. The carbon isotope abundance ratio was calculated using the international standard VPDB (Vienna Peedee Belemnite) to determine the δ^13^C value, with an analytical precision of ±0.2‰. Similarly, the nitrogen isotope abundance ratio was based on atmospheric nitrogen to calculate the δ^15^N value, with an analytical precision of ±0.25‰. Additional data on other macrobenthos in the Yellow Sea can be found in **Table S2** (*15*).

### Transmission electron microscopy

Gill tissues were fixed overnight using a solution containing 2.5% glutaraldehyde and 2% paraformaldehyde (PFA) in phosphate buffer (PBS). The tissue was washed in 0.1M PBS three times for 15 minutes each and then fixed with 1% osmium tetroxide (OsO4) for 1 hour, followed by additional wash using PBS. It was dehydrated through a methanol series (50%, 70%, 90%, and 100%, three times for 15 minutes each) and embedded in Epon 812 resin. Ultrathin sections (70 nm) were sliced using a Reichert ULTRACUT slicer (Austria) and stained with uranyl acetate and lead citrate double staining method (*42*) for 15 minutes each. Images were captured by a JEM 1200-EX (Japan) transmission electron microscopy (TEM) at an accelerating voltage of 80 kV.

### Fluorescence *in situ* hybridization (FISH) and histology

For FISH experiments, the symbiont-specific probe with Cy5-labeled (5’-TCCTCTATCACACTCTAGCTCAGCAGTATC-3’), sense probe with CY3-labeled (5’-GATACTGCTGAGCTAGAGTGTGATAGAGGA-3’), and bacterial universal probe EUB338 with CY5-labeled were designed based on the corresponding representative 16S rRNA gene (*43*). Primer-BLAST was used to evaluate the specificity of the designed probes, with reference of the NCBI non-redundant nucleotide sequence database (*44*). Gill tissues (fixed in PFA and preserved in pure methanol) were dehydrated in 100% methanol for 30 minutes each, embedded in paraffin, and then sectioned into 7 μm thick slices using a semiautomatic microtome (Leica, Germany). After removing paraffin with xylene and ethanol, the sections were rehydrated in a decreasing ethanol series (100%, 95%, 80%, and 70%) for 15 minutes each, followed by hybridization at 46°C with a hybridization buffer (work concentration: 5 μg/mL probe in 0.9 M NaCl, 0.02 M Tris-HCl, 0.01% sodium dodecyl sulfate and 30% Formamide) for 1 h. Following hybridization, the slides were washed in a washing buffer (0.1 M NaCl, 0.02 M Tris-HCl, 0.01% sodium dodecyl sulfate, and 5 mM EDTA) at 48°C for 5 minutes each, and subsequently, the cell nucleus and cell membrane were stained with 4′,6-diamidino-2-phenylindole (DAPI, Solabio) and Alexa Fluor 488 Conjugate Concanavalin-A (Invitrogen, CA, USA) for 10 minutes at room temperature, respectively. After washing using PBST (Tween-20: PBS=1: 1000), the slides were mounted with ProLong Diamond Antifade Mountant (Invitrogen). Images were captured using a ZEISS LSM900 or Andor Dragonfly 302 confocal laser scanning microscope. For hematoxylin and eosin (HE) staining, dewaxed tissue sections were stained according to standard protocols. Subsequently, sections were dehydrated and mounted with neutral balsam. Images were captured by a pathological section scanner (Leica SDPTOP HS6)

### Full-length amplicon sequencing

Genomic DNA was extracted from the whole or partial gill tissues (The whole gill divided into six parts) using the DNeasy Blood & Tissue Kit (Qiagen, Hilden, Germany), following the manufacturer’s protocol. Meanwhile, DNA was also extracted from approximately 0.5 g of ambient surface sediments (wet weight) by using the PowerSoil DNA Isolation Kit (Qiagen, Hilden, Germany). NanoDrop Lite (Thermo Scientific, USA) and 1% agarose gel electrophoresis were used to check the DNA quantity and quality, respectively. Full-length 16S rRNA gene of bacteria from sediments and *T. tokunagai* gill were also amplified by the primers 27F and 1492R (*45*). High fidelity (HiFi) reads were generated from the PacBio RS II platform by Novogene (Beijing, China) with CCS mode in gill samples and ambient sediment samples, respectively. QIIME version 2023.9.1 (*46*) was used to data process with the standard pipeline, containing quality control, Amplicon Sequence Variants (ASVs) / phylotypes table construction, and taxonomic classification. ChiPlot (https://www.chiplot.online) was used to visualize the relative abundance of the bacteria community in the gill and sediment.

### Metagenome sequencing

Genomic DNA intended for amplicon sequencing and the newly extracted DNA were both employed for metagenomic sequencing. The newly extracted genomic DNA was obtained from gill and whole tissue using the SDS method. The library was constructed by randomly fragmenting the DNA into approximately 350 bp reads. Following library construction, sequencing was conducted in paired-end 150 bp mode on an Illumina NovaSeq 6000 platform (Tianjin, China). Simultaneously, the long quality-checked DNA was sent to Novogene (Tianjin, China) for library preparation and sequencing. For Oxford Nanopore Technologies (ONT) sequencing, the library was generated using the SQK-LSK109 kit (Oxford Nanopore Technologies, UK) in accordance with the manufacturer’s guidelines. Long raw reads were generated through basecalling with Guppy version 6.1.7 (*47*).

### Metatranscriptome sequencing

Total RNA was extracted using TRIzol reagent (Invitrogen, CA, USA) with the guidance of the manufacturer’s protocol. RNA integrity and quantity were measured using the Bioanalyzer 5400 system (Agilent Technologies, CA, USA). cDNA was obtained by removing the prokaryotic and eukaryotic ribosomal RNA (rRNA of Animal, G-Bacteria, and Plant) from the total RNA using the TIANSeq rRNA Depletion Kit for the construction of a meta-transcriptomic library. The nucleic acid for metagenome and meta-transcriptome sequencing was subjected to NovaSeq 6000 system (Illumina) at Novogene (Tianjin, China) with paired-end mode and a read length of 150bp, and the ONT library was sequenced on the PromethION platform at Novogene.

### Mitochondrial genome assembly and annotation

Raw reads were trimmed to remove low-quality sequences and adapters using Trimmomatic v.0.39 (*48*) with the following parameters: ILLUMINACLIP: TruSeq3-PE-2.fa:2:30:10, LEADING:20, TRAILING:20, SLIDINGWINDOW:4:15, MINLEN:100. NOVOPlasty v.4.3.1 (*49*) were employed to construct the mitochondrial genomes with default settings (160M randomly selected reads per sample). Assembled mitochondrial genomes were annotated on the MITOS web server (*50*) with the default setting except “the genetic code: 5 invertebrates”.

### Host phylogenetic and population structure analysis

A total of 24 *cox1* gene sequences in the family Thyasiridae were downloaded from the NCBI, and three Lucinid sequences served as the outgroup in the phylogenetic analysis **(table S8)**. Then, these sequences were aligned using MUSCLE v.5.1 (*51*), and the ambiguous alignment was trimmed using Gblocks v.0.91b (*52*). The phylogenetic tree was constructed using the Maximum Likelihood (ML) method in MEGA-X v.10.2.2 (*53*) and 100 bootstraps, and the HKY + Γ model was suggested as the best model based on both AIC- and BIC-based methods.

To understand the population differentiation of *T. tokunagai* in the Yellow Sea, the DnaSP v.6 (*54*) and PopART v.1.7 (*55*) with the median-joining Network method were employed to investigate the haplotype network based on the alignment file. Furthermore, 13 protein-coding genes (PCGs) of 30 individuals were aligned separately using MAFFT v.7.515 (*56*) with the default parameter. Population structure analysis was performed based on the mitochondrial genome by STRUCTURE v.2.3.4 (*57*) with the settings of “*K*: from 2 to 7, 2,000,000 iterations, and 10% of burnin”. The most optimal *K* was determined using Structure Harvester (*58*) web server with the delta *K* method.

### Symbiont genome assembly, binning, and annotation

Short raw reads were trimmed using Trimmomatic version 0.39 (*48*) with the following settings: TruSeq3-PE-2.fa:2:30:10:8:true SLIDINGWINDOW:5:20 LEADING:3 TRAILING:3 MINLEN:36. Clean reads were assembled using Megahit version 1.2.9 (*59*) with default settings. The metagenome-assembled genomes (MAGs) were binned using MaxBin version 2.2.7 (*60*), with the cutoff of contig length from 1000 to 2000 for the optimization. To reduce the host contamination, we conducted a decontamination process using BlobTools version 1.1.1 (*61*) with default settings, and then sequences belonging to the phylum *Proteobacteria* were selected for downstream analyses. To obtain the circular-level genome of the symbiont, firstly, the ONT long reads were mapped to the highest quality MAGs; secondly, these long reads mapped were assembled using NextDenovo version 2.5.2 (*62, 63*); thirdly, clean reads from each sample (i.e. Illumina) were input into NextPolish version 1.4.1 (*64*) to polish the circular-level genome. Then, these genomes were evaluated for completeness and contamination using CheckM2 version 0.1.3 (*65*). GTDB-TK version 2.1.1 (*66*) was used to determine the taxonomy of symbionts at the genome level. The matrix of pairwise average nucleotide identity (ANI) of 30 MAGs was generated using FastANI version 1.34 (*17*). The Wilcoxon Rank-Sum test was employed to assess differences in ANI values across symbiont phylotypes. 16S rRNA genes and open reading frames (ORFs) of MAGs were predicted by Prokka version 1.14.6 (*67*) in the single genome mode. These predicted genes of the pangenome were searched against the NR database using BLASTp in DIAMOND version 2.0.15.153 (*68*) with an E-value cut-off of 1e^−5^. The results were further used for Gene Ontology annotation by Blast2GO version 6.0 (*69*). Meanwhile, Clusters of Orthologous Group 2020 (COG2020) (*70*) was adopted to classify the functional groups of genes in the pangenome. The genes of the pangenome were annotated using BlastKOALA (*71*) by searching against the KEGG database **(table S9)**.

### Spatial metabarcode sequencing

We used the Stereo-seq FFPE pipeline (BGI, China) to investigate the spatial distribution pattern between the two 16S phylotypes. The gills from six individuals were pre-fixed overnight with 4% PFA and then embedded in paraffin. Three continuous sections from the embedded block were cut at 10 μm thick. The second section was designated for chip loading to capture rRNA, while the remaining two sections were used in staining (either HE or FISH). The tissue section was adhered to the Stereo-seq chip (BGI, China) surface and incubated at 37□ for 3 minutes. The tissue sections were fixed in methanol and incubated at −20°C for 40 minutes prior to Stereo-seq library preparation. Where applicable, the same sections were stained with a nucleic acid dye (Thermo Fisher, Q10212), and imaging was conducted using a Stereo OR 100 microscopes in the FITC channel before *in situ* capture. After washing with 0.1X SSC buffer (Thermo, AM9770) supplemented with 0.05 U/ml RNase inhibitor (NEB, M0314L), the tissue sections placed on the chip were permeabilized using 0.1% pepsin (Sigma, P7000) in 0.01 M HCl buffer. They were incubated at 37°C for 5 minutes and subsequently washed again with the same 0.1X SSC buffer with RNase inhibitor. cDNA was synthesized on the chip using the FFPE MIX solution, consisting of 158 μL FFPE RT Buffer mix, 30 μL FFPE RT Enzyme mix, 10 μL FFPE RT Oligo, and 2 μL FFPE Dimer, at 42°C for 5 hours. The cDNA-containing chips then underwent treatment with the Prepare cDNA Release Mix (cDNA Release Enzyme and cDNA Release buffer) overnight at 55°C. The harvested cDNA was purified using VAHTSTM DNA Clean Beads (0.8X) and subsequently amplified in the amplification solution, which included 42 μL cDNA, 50 μL cDNA amplification mix, and 8 μL FFPE cDNA Primer Mix. The PCR protocol was as follows: initial denaturation at 95°C for 5 minutes, followed by 15 cycles of denaturation at 98°C for 20 seconds, annealing at 58°C for 20 seconds, and extension at 72°C for 3 minutes, concluding with a final extension at 72°C for 5 minutes. After quantification using the Qubit dsDNA HS kit, the cDNA product was utilized for library construction according to the guidelines of the Stereo-seq 16 Barcode Library Kit V1.0. Raw reads were retrieved in paired-end 75bp mode using MGI DNBSEQ-T7.

### Spatial metabarcode analyses

Fastq files were generated from an MGI DNBSEQ-T7 sequencer. CID and MID are contained in the read 1 (CID: 1-25 bp, MID: 26-35 bp), while the corresponding read 2 consists of the 16S rDNA sequences. The current SAW software version 8.0.2 (*72*) could not differentiate the two 16S phylotypes with the difference of a single base pair. To visualize the patterns, the pre-separation of reads containing the differentiated base pair fully matching to the full length 16S rRNA gene were adopted. In detail, the normal analysis of Stereo-seq reads were completed, with all the raw reads, ssDNA image, mask file, and reference. The STAR genome reference was prepared with the *de novo* assembled transcriptional profile and the 16S rRNA sequence (phylotype A). The produced *.bam* files were processed, with the region (16S rRNA sequence: 907-953) kept. The A-phylotype reads were extracted if they were uniquely mapped and completely matched to 16S rRNA (NH:i:1, MAPQ=255, CIGAR=47M). Meanwhile, the G-phylotype reads were extracted as the same way, with the substitution of a single base pair (A → G). Then, two independent analyses of them were run using SAW pipeline. The count of mapping reads spatially was shown as the two-dimensional locations and the corresponding mapping number (the tab-delimited file with three columns). Given the cellular size (about 10 μm), these counts were aggregated into bin 20 (20×20 DNBs). Bin 20 spots were considered valid only if they had at least 180 reads. The ratio of 16S phylotype (A or G) was then calculated, with the frequency histogram shown (**fig. S10**). The dominant phylotypes in bins were defined if the ratio was above 95%. The result was visualized using matplotlib in Python.

### Phylotype decomposition, phylogenomic analysis of symbiont, and comparison

Pangenome was generated using PanPhlAn version 3.1 (*73*) and in-built scripts of StrainPanDA (*74*). To decompose the diversity of symbiont at the high resolution, StrainPanDA (*74*) was adopted based on the newly constructed pangenome of 30 MAGs and the full set of clean reads. The phylogenetic positions of the two symbiont phylotypes were reconstructed among known chemosynthetic bacteria, utilizing 57 published SOB genomes and the *B. azoricus* MOB symbiont as an outgroup. The phylogenomic analysis was conducted using VEHoP version 1.0 (*75*) to determine their positions at the phylogenomic level. The phylogenomic tree was visualized using tvBOT (*76*). To differentiate between the two symbiont phylotypes, a functional information comparison analysis was performed using METABOLIC□G version 4.0 (*77*), based on the strain decomposition results.

### Transcriptional profile assembly and quantification

The raw reads were filtered using trimmomatic version 0.39 with the settings (TruSeq3-PE-2.fa:2:30:10:8:true SLIDINGWINDOW:5:20 LEADING:3 TRAILING:3 MINLEN:75). Then, qualified reads were mapped to pangenome using Bowtie version 2.3.5 (*78*), resulting in the symbiont-derived reads and symbiont-free reads. The symbiotic-free reads were subjected to Trinity version 2.13.2 for de novo assembly (*79*). To prepare for spatial metabarcoding analysis, the host transcripts were first purified using BlobTools version 1.1.1 (*61*) to remove contaminated contigs, and subsequently assessed for coding potential using TransDecoder version 5.7.1 (*80*). Salmon version 1.9.0 (*81*) was performed to quantify the gene expression levels of symbiont.

### Carbon fixation rate measurements

Radiotracer assays were used to determine the rate of dissolved inorganic carbon (DIC) assimilation by introducing a ^14^C-labeled DIC tracer to the homogenized gill in sterilizing seawater and quantifying the amount of ^14^C incorporated into total organic carbon (TOC) (*82*). Gill tissues dissected from fresh samples were homogenized and immediately stored in 20% glycerol in sterile seawater, then slowly thawed on ice prior to downstream analyses. The experimental design details are as follows (**Fig. 5B**): 1) to minimize the impact of varying symbiont numbers on our results, we collected samples from four sites, and four individuals per site; 2) to create a uniform starting point, samples from each site were mixed, and the resulting mixture was divided into sixteen replicates, and these replicates were then assigned to one of four incubation groups at different temperatures (5□, 12□, 20□, and 28□), and incubated in the dark for 46-47 hours; 3) Within each temperature group, we prepared four replicates of gill tissue samples: three experimental samples and one negative control. These samples were placed into 10 ml serum vials, ensuring no headspace, and sealed with sterile PTFE septa and aluminum caps. After that, 100 µL of ^14^C-DIC solution (~4 × 10^4^ Becquerel, Bq) was injected into each serum vial through the stopper by displacing the same volume of water. Before injecting the ^14^C-DIC tracer, the microorganisms of negative controls were killed by adding 0.5 mL of 100% trichloroacetic acid. Microorganisms of the experimental group were removed with the addition of 0.5 mL and filtered onto 0.2 μm GSWP membranes (polyethersulfone, Millipore) after incubation. The filters were rinsed with 35 ‰ sodium chloride (NaCl) solution (*83*) and transferred into 7 mL scintillation vials containing a 6 mL scintillation cocktail (Ultima Gold ™ Cocktail, PerkinElmer). The radioactivity of the filters was determined using a Tri-Carb 3110TR liquid scintillation counter (*84*). The turnover rate constant (*k_n_*) of DIC was calculated using the equation No. (1), and the assimilation rate (Ass-rate) of DIC was calculated using the equation No. (2):

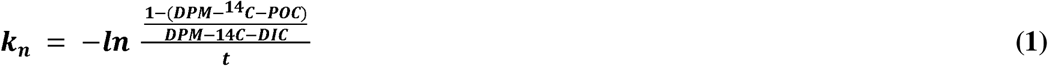

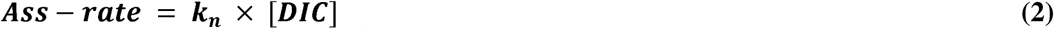

*k_n_*: turnover rate constant = day^−1^.

*Ass-rate*: the DIC assimilation rate (µmol·L^−1^·day^−1^).

*DPM-^14^C-POC*: the radioactivity on the filter.

*DPM-^14^C-DIC*: the total activity of the added DIC tracer.

*t*: the incubation time (day).

*[DIC]*: the DIC concentration (mmol·L^−1^) ranged from 2.019 to 2.229 used in this study.

### Estimation of the carbon fixation flux of *Thyasira* in the Yellow Sea

We assumed the spatial variation in bottom-layer temperature and thyasirid density in the Yellow Sea to be a result of continuity, though the geological, chemical, and biological processes may also influence this. To avoid over-estimation, the total region for flux estimation was restricted by the maximum region from 139 sampling sites from seven cruises mentioned above (**Fig. 1A**). The bottom water temperature data for this study were derived from the mean annual values for the period 2002-2017 of the World Ocean Atlas 2018 (*85*). Meanwhile, biological distribution data of *Thyasira gouldii* complex (including *T. tokunagai*) were obtained from the Ocean Biodiversity Information System (OBIS; https://obis.org/) and the NBN Atlas (https://nbnatlas.org/). The Python package pykrige.ok version 1.7.2 was employed for the kriging interpolation among sites, predicting the spatial pattern of species density and bottom-layer temperature with a resolution of 0.005 degrees (division unit).

The estimation details of total annual carbon fixation flux in the Yellow Sea contributed by this clam are as follows: we measured the DIC assimilation rate at different temperature, while fitted the curve to reconstruct the relationships between turnover rate constant (*k_n_*) and bottom temperature. Based on this relationship, we used the openly available bottom water temperature of the World Ocean Atlas 2018 to predict the turnover rate constant. Then, the DIC assimilation rate was obtained by the DIC concentration multiplied by the turnover rate constant (*k_n_*). Annual carbon fixation flux (Ann-Flux) in each site was calculated using the equation No. (3), mainly incorporating the following parameters: (I) temperature-dependent rate constants (k) from radiocarbon assimilation experiments on *T. tokunagai* (calculated by the equation No. (4); Wilcoxon rank-sum test for site differences at same temperature), (II) interpolated *T. tokunagai* density, (III) interpolated temperature, and (IV) DIC concentration. The total annual carbon fixation flux is the sum of each division unit.

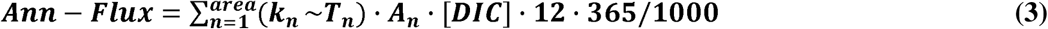

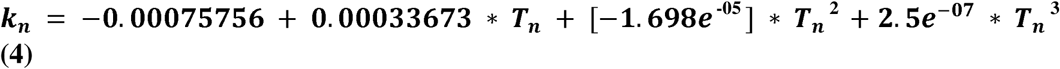

*Ann-Flux*: annual carbon fixation flux (g C · yr^−1^).

*k_n_~T_n_*: the relationship between rate constant and temperature.

*T_n_*: the predicted spatial temperature via krige exploitation (□).

*A_n_*: the predicted spatial abundance of *Thyasira tokunagai* via krige exploitation (m^2^ individual^−1^).

*[DIC]*: the concentration ranged from 2.019 and 2.229 (mmol·L^−1^).

12: the molar mass of carbon (g·mol^−1^).

365: assuming the total number of days in a year (day).

## Acknowledgments

This work was financially supported by the National Natural Science Foundation of China (42476135), Science and Technology Innovation Project of Laoshan Laboratory (LSKJ202203104 and 2022LSL050104-6), the Fundamental Research Funds for the Central Universities (202172002 and 202241002), and the Young Taishan Scholars Program of Shandong Province (tsqn202103036). Data and samples were collected onboard R/V “*Lanhai 101*” implementing the open research cruises NORC2020-01 and NORC2021-01 supported by the NSFC Ship Time Sharing Project (project numbers: 41949901 and 42049901). We thank Suzanne C. Dufour (Memorial University of Newfoundland) for her constructive comments during the preparation of this paper.

## Author contribution

JS conceived the project. ML collected the samples, extracted the nucleic acid, and performed most of the experiments and bioinformatic analysis. YL performed the strain decomposition and the carbon fixation estimations. ML and YL drafted the original manuscript. SM estimated the DIC assimilation rate of the symbiont. CC identified the specimens. YL, MS, CC, YL, XL, GCZ, WZ, and JS contributed to the writing and editing of the manuscript.

## Data and materials availability

The data of this study were deposited in the NCBI database under the BioProject ID PRJNA995037 and PRJNA1016492. All data are available in the main text or the supplementary materials.

## Notes

### Competing Interest Statement

The authors have declared no competing interest.

### Summary of Updates

Remove the former priority effect conclusion; Include a spatial metabarcoding analysis; Include more results on the radiocarbon tracing and spatial modelling analyses.

