## Supplementary information for "Shallow-water chemosymbiotic clams are a globally significant and previously overlooked carbon sink"

Menggong Li *et al.*

Guang-Chao Zhuang,

**This PDF file includes:**

Figs. S1 to S10

Legends for tables S1 to S9

**Other Supplementary Materials for this manuscript include the following:**

Tables S1 to S9

**Materials and Methods**

**Sampling description**

In detail, 1) samples for genomic DNA extraction were stored at 70 % ethanol or frozen at -80 ℃; 2) samples for fluorescence *in situ* hybridization (FISH) were initially fixed in 4 % paraformaldehyde (PFA) at 4 ℃ overnight and then transferred to 100 % methanol at -20℃; 3) samples for RNA extraction were preserved at RNA stabilization solution (Thermo Fisher Scientific Inc., Waltham, USA) at 4 ℃ overnight and long-term preserved at -80℃; 4) gill tissue for radioactive isotope tracing experiment were freshly dissected, homogenized at 60 % sterile glycerol solution, and flash-frozen at -80 ℃; 5) samples for transmission electron microscope (TEM) were stored at the mixed solution (2.5 % glutaraldehyde in phosphate buffer and 2 % PFA) at 4℃; and 6) samples for stable isotopic analysis were at dissected for the removal of shell and then kept at -20 ℃. In addition, six ambient surface sediment samples from two sites (NYS1 and NYS3) were collected and immediately frozen at -20 ℃ for the environmental microbial community analysis.

**Phylogenetic analysis of symbiont based on 16S rRNA gene**

Similarly, a total of sixteen 16S rRNA gene sequences were downloaded, and the 16S rRNA gene tree of symbiont was constructed with the *Escherichia coli* 16S rRNA gene as the outgroup. The alignment analysis and phylogenetic method were similar to the *cox1* gene analysis. The best model was the HKY +Γ +I model and 100 bootstrap was also used to evaluate the confidence level.

**Results**

**Clone library construction (Supplementary Note 1)**

To ensure that the difference of one nucleotide between the two oligotypes is not caused by sequencing error, the DNA of two representative samples (i.e., “NYS3_4” and “NYS3_5”) were used to construct the clone library. Firstly, to obtain the gene target fragment, the full-length 16S rRNA genes were amplified following the main text abovementioned protocol, and the PCR product was further purified by using the E.Z.N.A. Cycle Pure Kit (Omega Bio-tek, Inc). Then, a total of 70 ng target fragments were inserted into the vectors. Additionally, all experimental procedures were followed by pClone007 Blunt Simple Vector Kit (Tsingke, China) with standard protocol. A total of 23 valid single clones were selected randomly for Sanger sequencing by BGI (Qingdao, China) from both ends. Raw reads of full-length 16S rRNA genes were assembled to a consensus sequence with the primers removed. Collectively, a total of 23 nearly full-length monoclonal sequences were obtained: 11 *Sedimenticola* sp. (ex *Thyasira tokunagai*) phylotype G, 1 strain of *Spirochaeta*_2 genus from the NYS3_4 sample, 8 *Sedimenticola* sp. (ex *Thyasira tokunagai*) phylotype G, and 3 strains of the genus *Spirochaeta*_2 from the NYS3_5 sample.

**Fig. S1.**

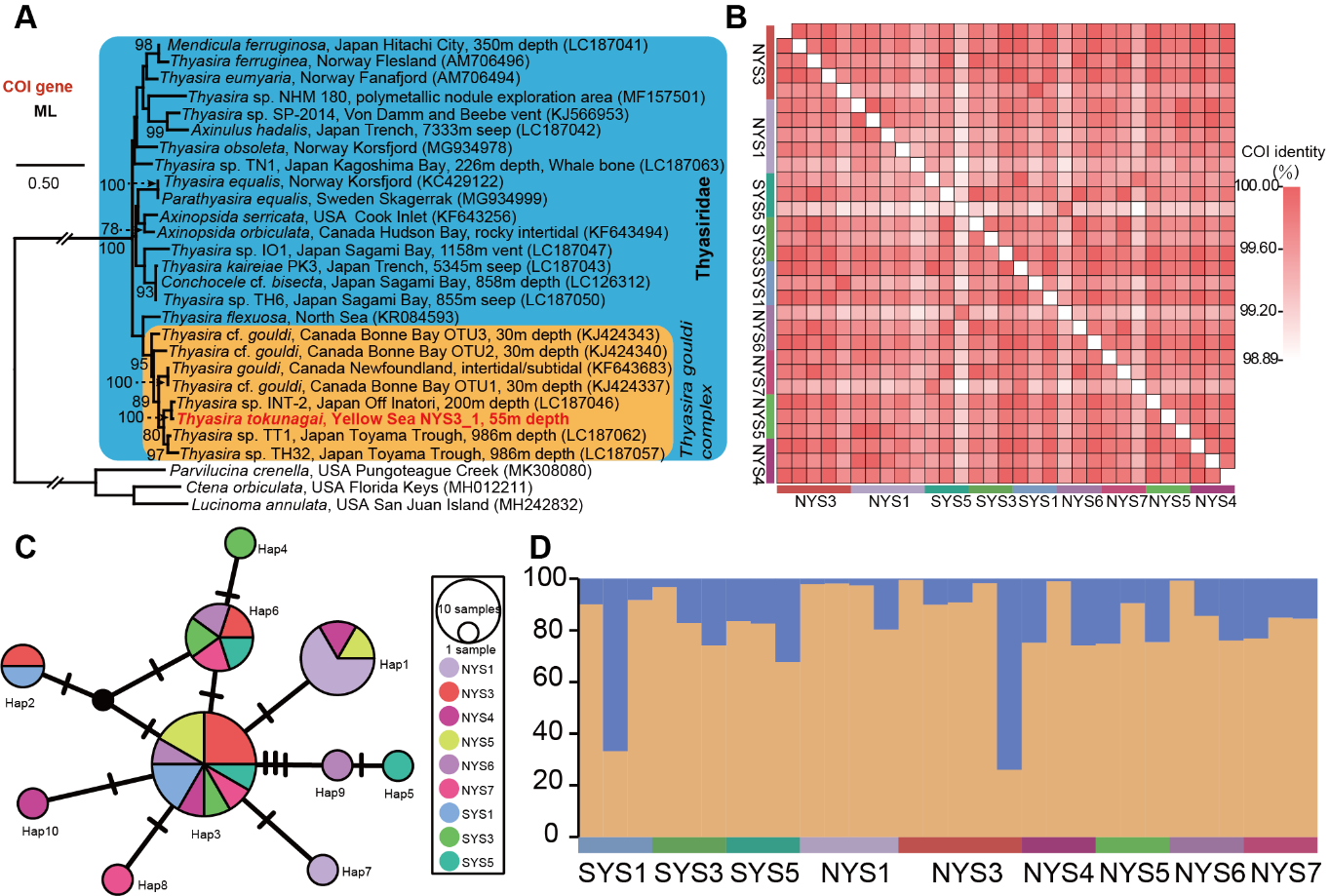

**Supplementary Figure S1: Phylogenetic and population analyses of *T. tokunagai* indicate a well-mixed population structure.** (A) Maximum-likelihood tree based on the partial *cox1* gene sequence of *T. tokunagai*. The representative *cox1* sequence of *T. tokunagai* is highlighted in red. This tree was constructed using the HKY +Γ model with 100 bootstraps. The scale bar indicates 0.50 substitutions per site. Values below 50 are hidden. (B) The pairwise similarity was performed based on the partial *cox1* gene sequences of *T. tokunagai* from 31 individuals located at a total of nine sites in the Yellow Sea. (C) The haplotype network of the *cox1* gene from 31 individuals with the median-joining network method showed the star-burst type, indicating there is population undifferentiation. (D) Host population structure analysis based on the 13 protein coding genes of the mitochondrial genome of 30 individuals from nine sampling sites in the Yellow Sea. This optimal delta *K* value of 2 was used in the host population structure analysis.

**Fig. S2.**

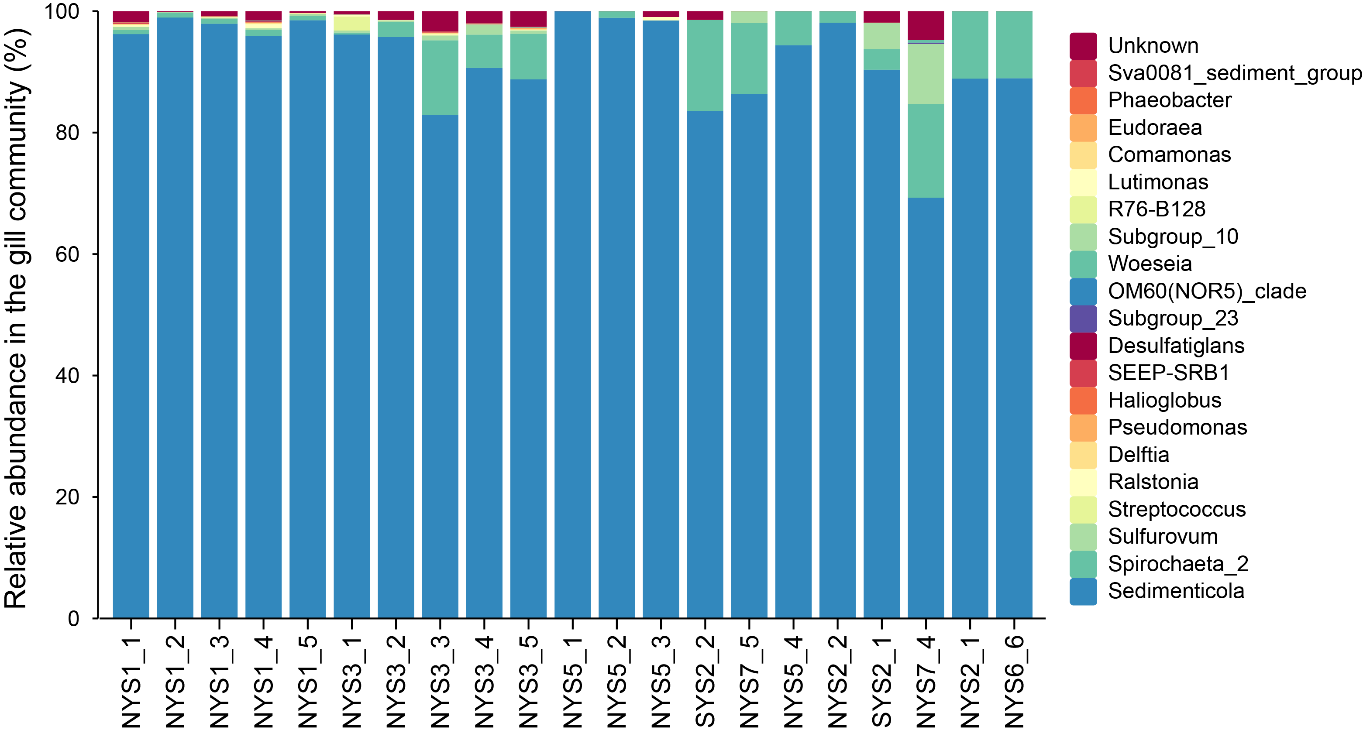

**Supplementary Figure S2: Genus-level bacterial community composition of gill tissue samples based on full-length 16S rRNA genes.** The results showed that the symbiont community was dominated by bacteria belonging to the *Sedimenticola* genus.

**Fig. S3.**

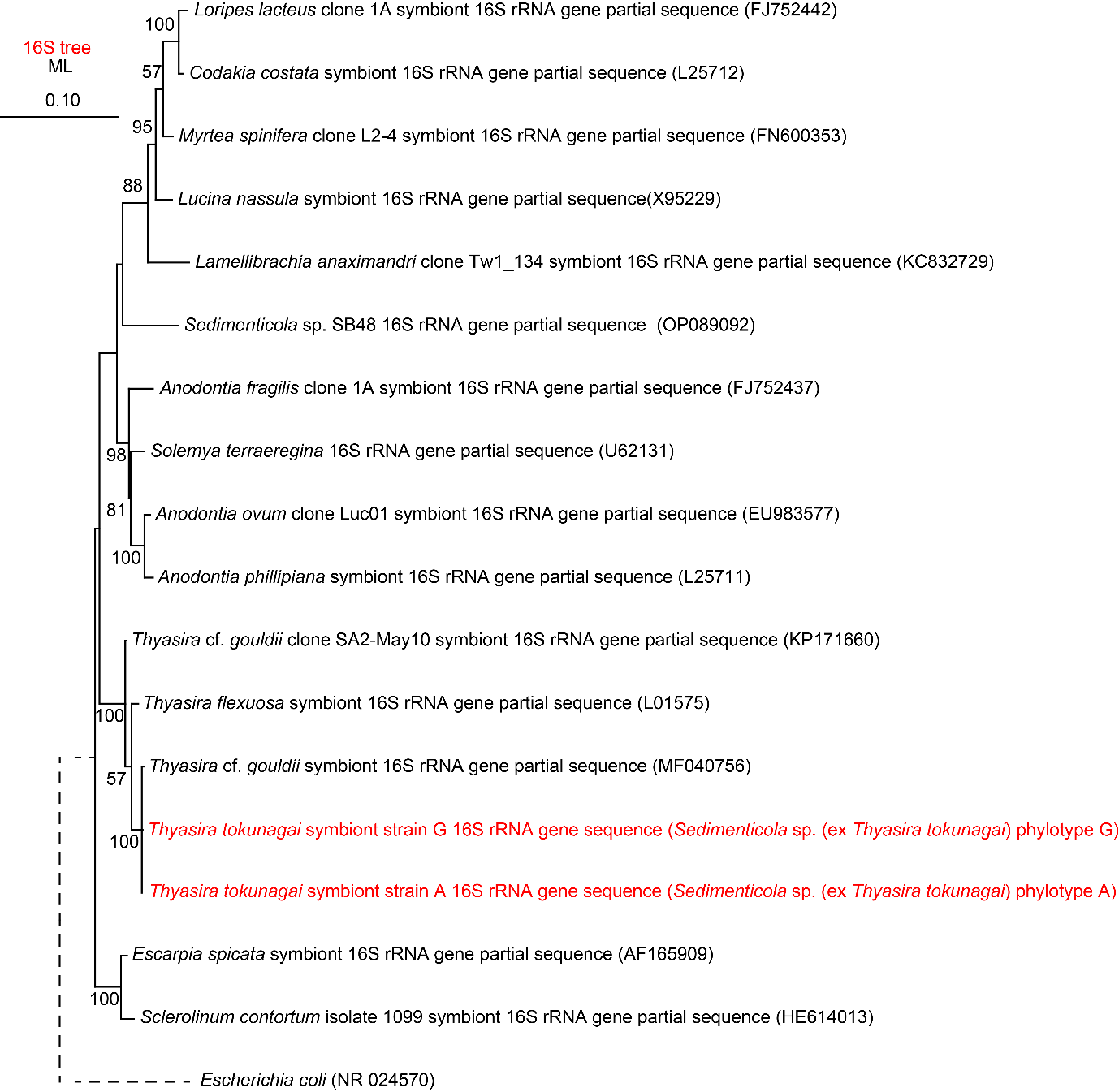

**Supplementary Figure S3: Phylogenetic analysis based on the full-length 16S rRNA gene sequences of symbiont confirmed the closely relative *Thyasira* cf. *gouldii*.** This tree was constructed based on HKY +Γ +I model with 100 bootstrap values. The scale bar indicates 0.10 substitutions per site. Values below 50 are hidden.

**Fig. S4.**

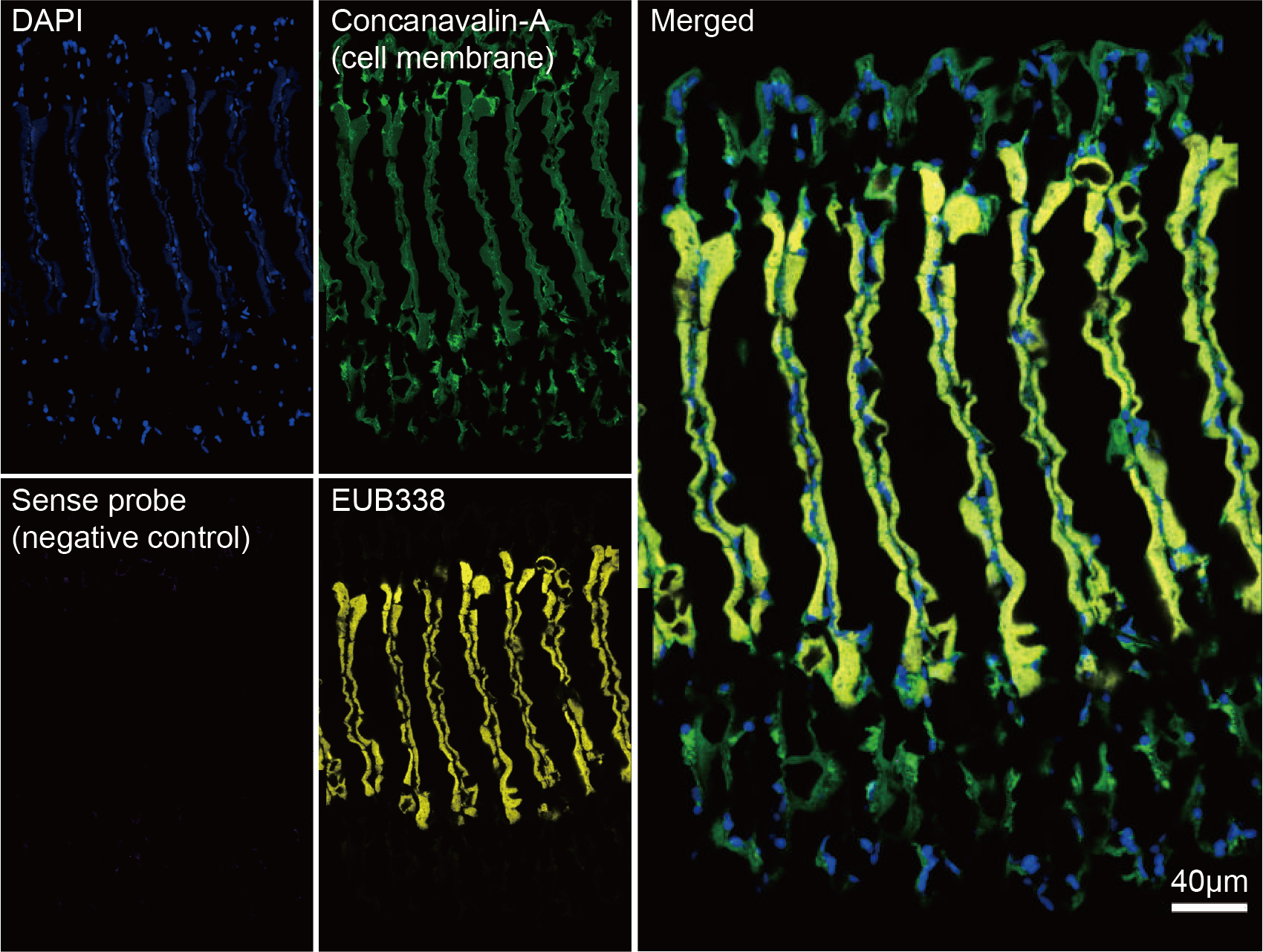

**Supplementary Figure S4: Fluorescence *in situ* hybridization based on the sense probe of symbiont showing the negative control signal in the gill tissue.** All cell nuclei were stained with DAPI, the cell membrane was stained by concanavalin-A, and the bacterium were hybridized by a universal probe of EUB338 with CY5-labeled.

**Fig. S5.**

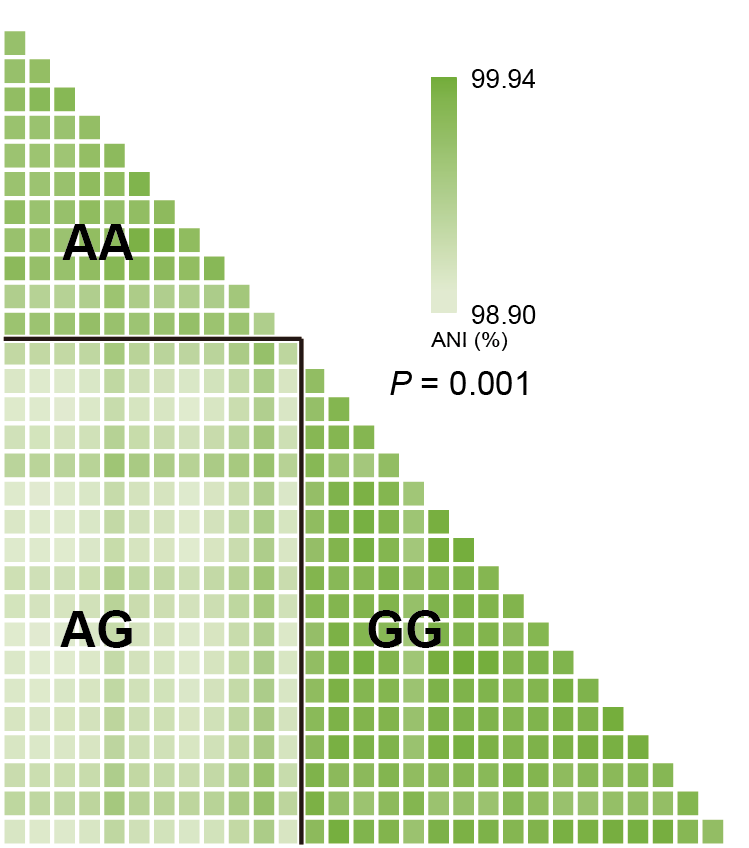

**Supplementary Figure S5:** **Average nucleotide identity (ANI) matrix of thirty symbiont population.** The results showed that we could detect genomic differences (*P* = 0.001) between the symbiont populations. Here, each host was treated as a symbiont population. "A" represented the sample was dominated by phylotype A, and "AG" represented the pairwise ANI value between two symbiont populations. Other labels followed the same convention.

**Fig. S6.**

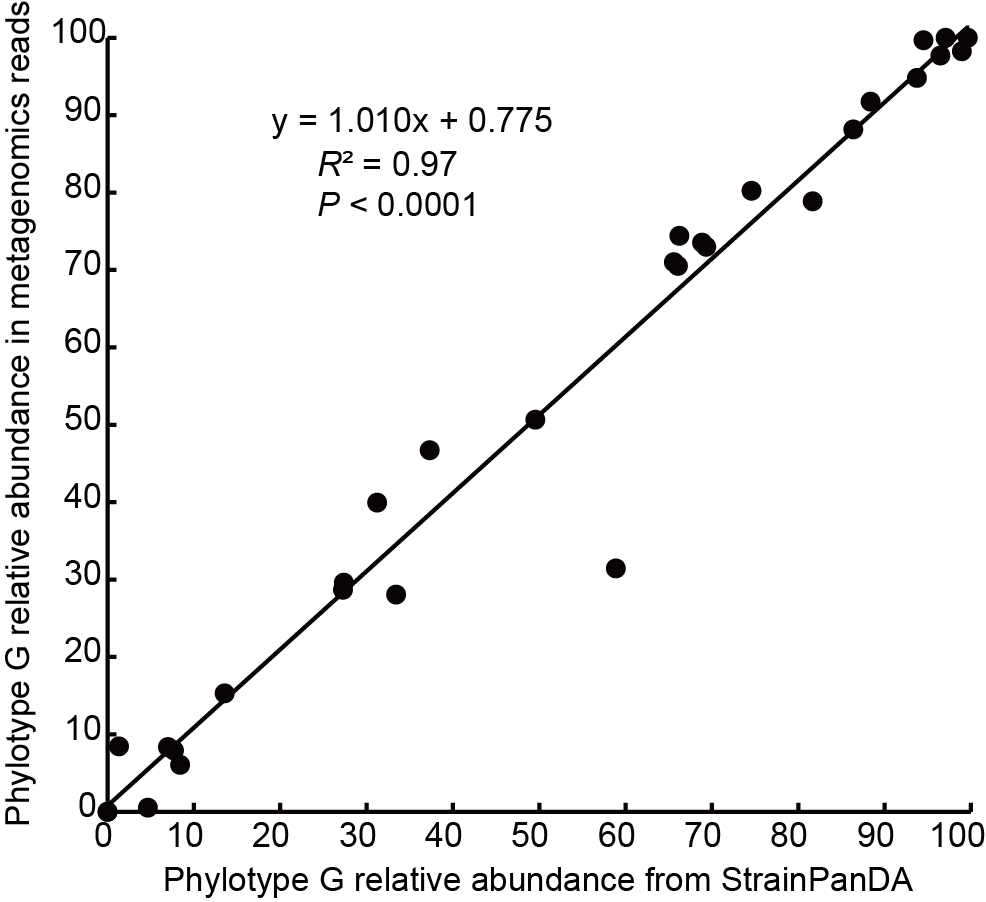

**Supplementary Figure S6: Correlation between phylotype G abundance and strain decomposition.** The correlation related to phylotype abundance in genome level between StrainPanDA and 16S rRNA gene relative abundance in each dataset with phylotype G as a quantitative (*R*^2^ = 0.97; *P* < 0.0001).

**Fig. S7.**

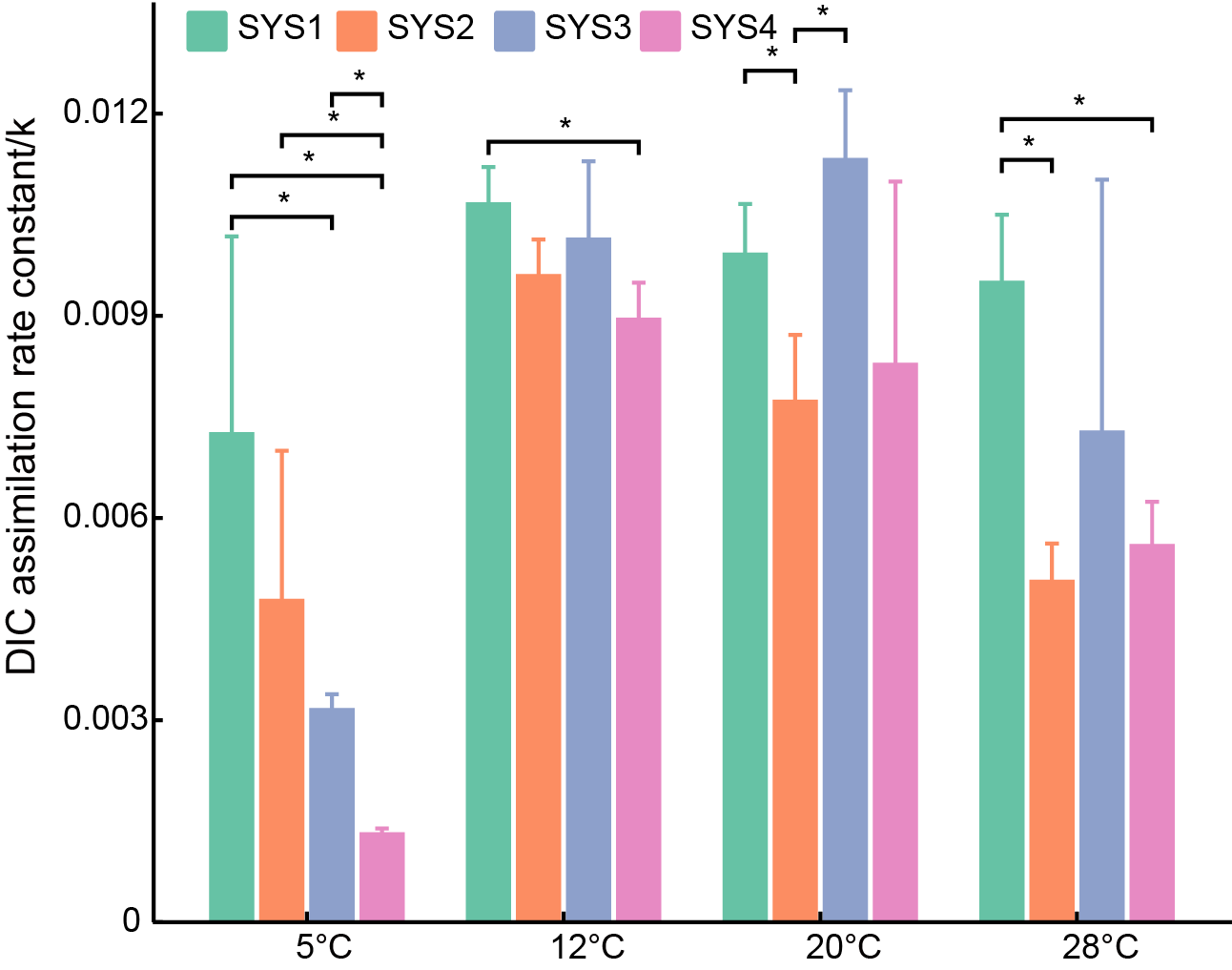

**Supplementary Figure S7: Temperature-related DIC assimilation rate constant at four sites in Yellow Sea.** Wilcoxon signed-rank tests revealed significant differences in DIC assimilation rate constants (k) at the same temperature, but only between certain Yellow Sea sites (SYS1-SYS4). As shown in the bar graph, significant differences *P* < 0.05, indicated by asterisks) were observed primarily at 5℃.

**Fig. S8.**

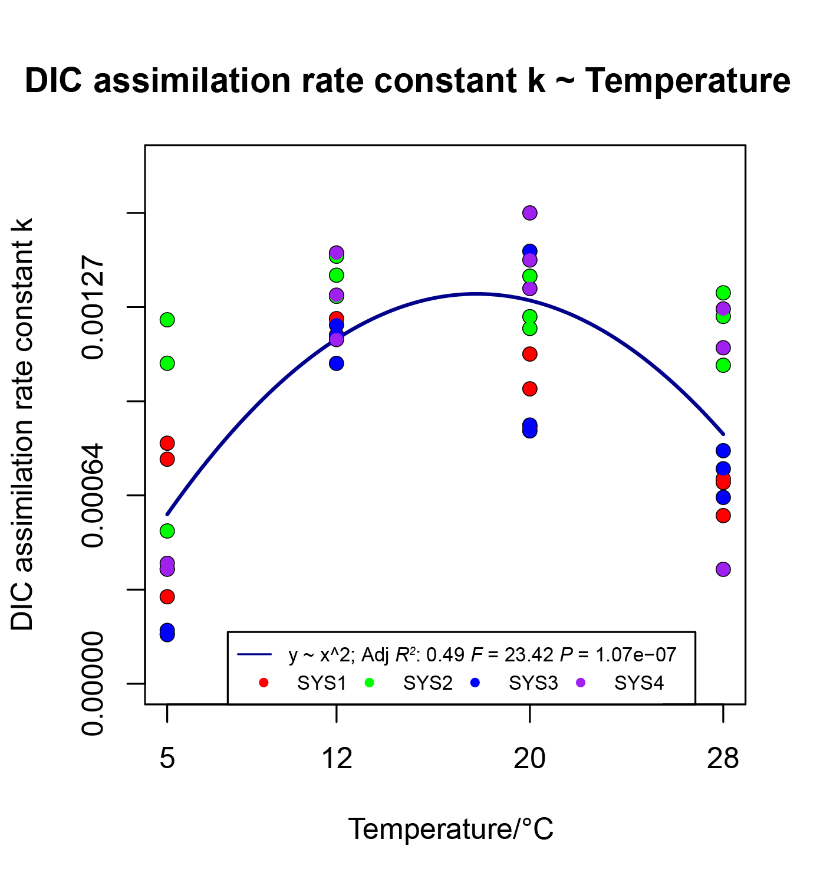

**Supplementary Figure S8: Carbon fixation rate constants at different temperatures (5°C, 12°C, 20°C, 28°C) were fitted using a binomial equation.** The equation as follows:

y = -0.00012398 + 0.00016214*x + -4.56e-06*x^2

**Fig. S9.**

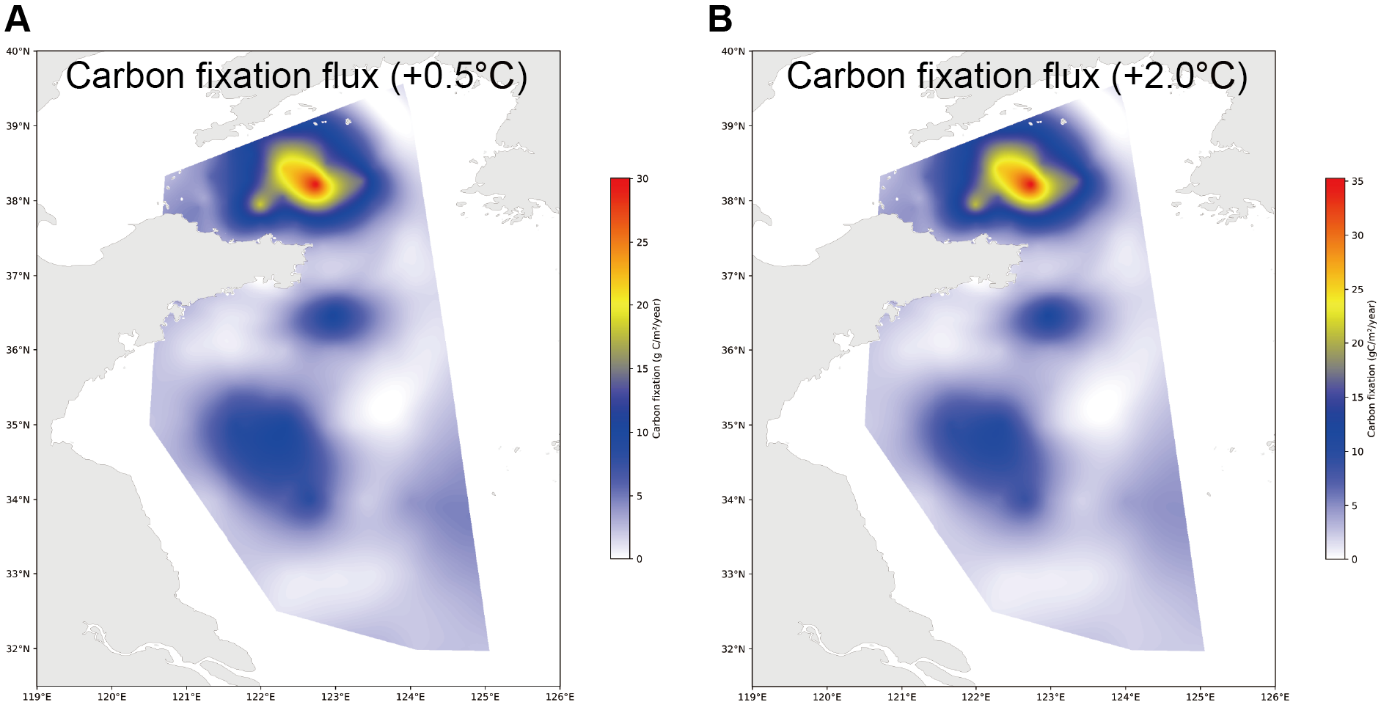

**Supplementary Figure S9: Estimation of annual carbon fixation flux when the bottom water temperature rises by 0.5℃ (A) and 2℃ (B) under the global warming environment.**

**Fig. S10.**

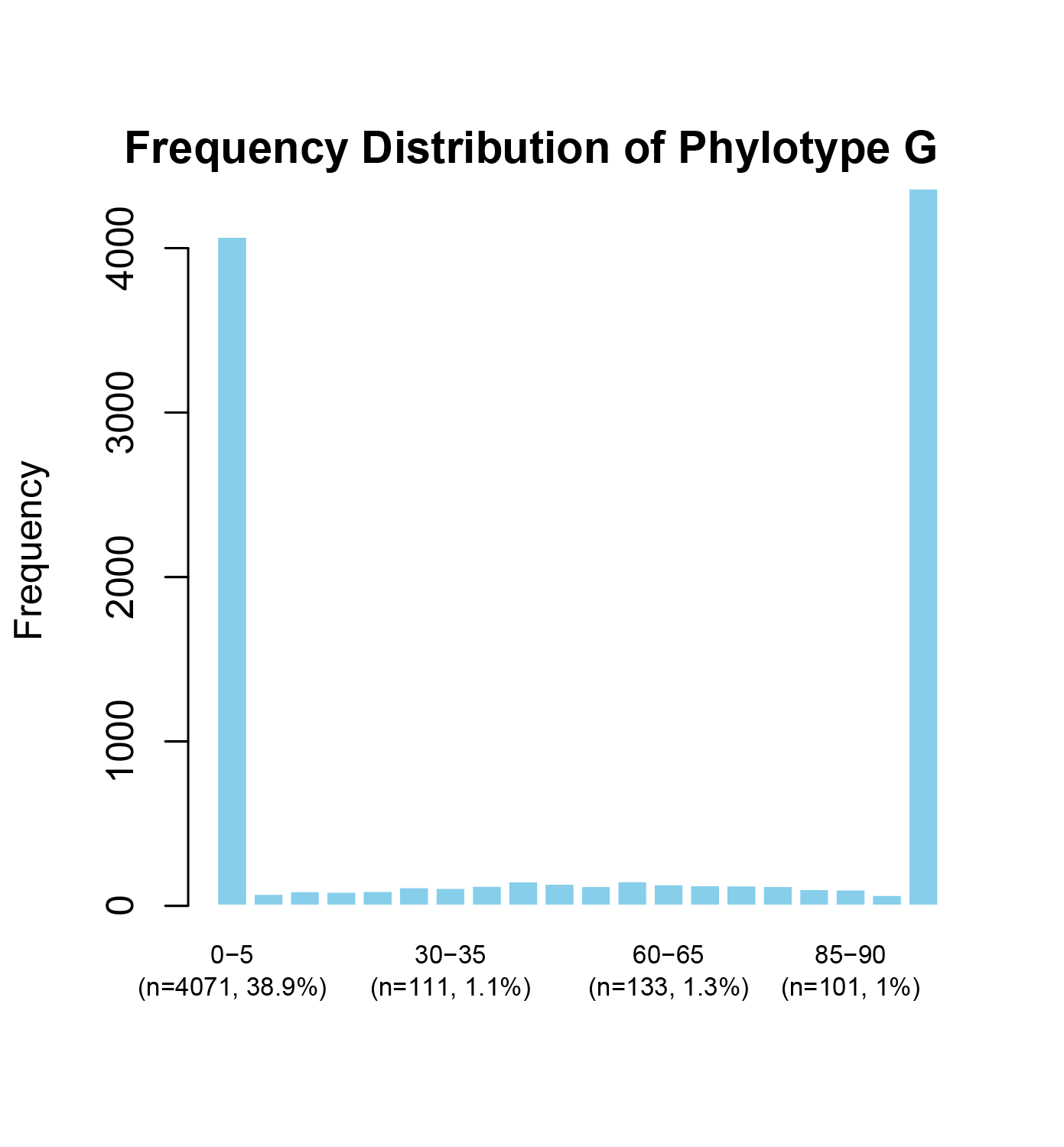

**Supplementary Figure S10: The ratio of 16S phylotypes (A or G) were then calculated in spatial metabarcoding analysis, with the frequency histogram shown.** Here, we statistic the distribution frequency of phylotype G ratio present in the total bin 20.

**Captains of Supplementary Tables:**

**Supplementary Table S1: Sampling information.**

**Supplementary Table S2: Stable isotopic data of samples used in this study from the Yellow Sea.**

**Supplementary Table S3: Statistics of *Thyasira tokunagai* symbiont genome assembly, and binning combining long-read and short-read sequencing.**

**Supplementary Table S4: Average nucleotide identity (ANI) matrix of thirty symbiont population.**

**Supplementary Table S5: Difference of functional coding genes between two symbiont phylotypes.**

**Supplementary Table S6: Metabolism capacity of symbiont about Amnio acid, vitamin, and cofactor biosynthesis.**

**Supplementary Table S7: Measurement records of DIC assimilation rate using radioactive carbon tracing.**

**Supplementary Table S8: Information about the cox1 gene sequence in constructing phylogenetic tree of *Thyasira tokunagai*.**

**Supplementary Table S9: Annotation information of functional coding genes of pangenome inferred from thirty symbiont genomes.**
